# Bacterial Mercury Resistance Reveals a Robust Species-Structured Human Antimicrobial Mobilome

**DOI:** 10.64898/2026.06.26.734528

**Authors:** Carolina López-López, Aida Ripoll, Antonio Sánchez Valenzuela, Sabrina Llop Pérez, Maria-Jose Lopez-Espinosa, Mónica Guxens, Mariona Bustamante, Amaia Irizar, Ziortza Barroeta, Lourdes Cirugeda, Fernando Baquero, Val F. Lanza, Teresa M. Coque

## Abstract

Antimicrobial resistance (AMR) is increasingly recognised as an ecological and evolutionary phenomenon that extends beyond clinical environments. Despite the predominant focus on antibiotics, bacterial metal resistance genes are among the oldest and most widespread adaptive antimicrobial systems, yet their distribution within human-associated microbial communities remains poorly characterised. Here, we investigated the occurrence of mercury (Hg) resistance and characterised the diversity of opportunistic pathogen species carrying *mer* genes, as well as their genetic contexts, and associated metal and antibiotic resistance determinants in children from a birth cohort with relatively high Hg exposure. Fecal samples from 234 children aged 4–8 years were analysed using culture-based screening, whole-genome sequencing, and comparative genomic analyses of resistance determinants and associated mobile genetic elements (MGEs). Hg-resistance was detected in 79.7% of samples, with Hg^R^ Enterobacterales isolated from 57% of children. Hair mercury concentrations were not associated with carriage. Sequencing revealed a phylogenetically diverse collection dominated by *Escherichia* spp. (61%). Hg resistance was mediated by 79 *mer* operons primarily associated with Tn*21*, Tn*1696*, and Tn*505*3 families circulating on chromosomes and a highly diverse plasmidome. Plasmids carrying *mer* operons were highly related to others previously reported in foodborne, animal, and clinical Enterobacterales. Metal resistance determinants exhibited strong taxonomic structuring, with *Escherichia* enriched in iron-uptake systems and siderophores whereas non-*Escherichia* taxa carried multiple metal resistance operons. These findings indicate that Hg-resistance is shaped by ecological interactions and MGEs, becoming partially decoupled from contemporary Hg-exposure and bacterial community composition. The human gut therefore serves as an important reservoir linking environmental metal resistance to the broader evolution of AMR and provides insight into the baseline resistome and plasmidome of human populations.

## Introduction

Antimicrobial resistance (AMR) is a global health challenge with far-reaching consequences for human, animal, and environmental health at the individual and ecosystem levels, as well as for economies and progress toward the United Nations Sustainable Development Goals ^1,2^. While research and policy efforts have primarily focused on antibiotics, resistance to heavy metals (HMs) has received comparatively little attention. Anthropogenic activities since the Industrial Revolution have led to the pervasive release of HMs into soils, waters, and the atmosphere through their widespread use in industry, agriculture, and biomedicine. In addition to threatening human and environmental health, and food safety, HM pollution exerts selective pressures on microbial communities that could drive the co-selection of antibiotic resistance ^3–6^. In parallel, metal-based antimicrobials represent a rapidly expanding research field ^7^, while the use of metals in industrial, agri-food, and household applications continues to grow, with global sales of biocidal agents including metals, projected to surpass those of antibiotics by 2030 ^8^. These numbers suggest that metal-associated selective pressures are likely to remain an increasingly important component of microbial evolution in the coming decades.

Among HMs, mercury (Hg) poses a particular threat to ecosystems and human health owing to its high toxicity and the dramatic increase in anthropogenic emissions above natural background levels ^9–11^. According to the latest global Hg assessment by the United Nations Environment Programme, environmental Hg burdens have risen by approximately 450% relative to natural levels, driving its accumulation across aquatic and terrestrial food webs ^6,9,10^. Present-day contamination is dominated by historical and ongoing emissions from artisanal and small-scale gold mining, together with continuing releases from industrial activities, waste management, and energy production ^9,12,13^. Importantly, the widespread use of mercurials in healthcare since the 18th century, including in disinfectants, antiseptics, dental amalgams, and medical devices ^13–15^, has been associated with the selection of Hg^R^ bacteria and the co-selection of antibiotic resistance ^15,16^.

Microbes play a central role in the global Hg cycle through the transformation, detoxification, and mobilisation of Hg compounds ^6,17^. These processes are primarily mediated by bacterial *mer* operons, which encode Hg detoxification functions and have undergone extensive horizontal transfer across taxa ^17–19^, including colonizing opportunistic pathogens (COPs) categorised as “antibiotic resistance threats” by the World Health Organization ^20,21^. The *mer* operons are frequently embedded within mobile genetic elements (MGEs) that also carry antibiotic resistance genes (ARGs), providing a genetic basis for the linkage between Hg resistance and antibiotic resistance ^3–5^.

Despite well-established Hg exposure pathways and health impacts ^11^, the ecological and evolutionary processes governing the persistence of Hg resistance in human-associated microbial communities remain poorly understood ^18^. In particular, it remains unclear whether the distribution of Hg resistance primarily reflects contemporary host exposure or the long-term ecological and evolutionary dynamics of mobile Hg^R^ determinants. Resolving this question is important not only for understanding Hg resistance itself, but also for clarifying how adaptive antimicrobial traits persist and circulate across microbial communities and ecological compartments. Europe, and particularly Mediterranean countries, provide a natural geographical setting for addressing these questions owing to their relatively high environmental Hg burden, shaped by historical, geological, industrial, and climatic factors, as well as fish dietary habits ^12,22^. Children provide a particularly informative biological framework for addressing these questions because their developing microbial communities can harbour greater phylogenetic and resistome diversity while lifetime antibiotic exposure remains limited ^23^.

Here, we investigated Hg^R^ bacteria in stool samples from children participating in the INMA cohort (*<u>IN</u>fancia y <u>M</u>edio <u>A</u>mbiente*, Environment and Childhood), a Spanish multicentre birth cohort characterised by relatively high prenatal and early-life Hg exposure, driven primarily by maternal fish consumption and sociodemographic factors ^24–27^. We combine phenotypic assays, genomic analyses, and MGE profiling to characterise the prevalence, genetic architecture, and ecological distribution of Hg resistance determinants in human gut bacterial communities, with a particular focus on Enterobacterales, which constitute major reservoirs of antibiotic resistance and frequent carriers of Tn*3*-family Hg transposons ^20,28,29^. We further investigate how Hg resistance determinants are embedded within, and potentially shaped by, a robust species-structured antimicrobial mobilome.

## Results

### Hg^R^ bacterial carriage and Hg exposure in the INMA cohort

We analysed 234 faecal samples from healthy children aged 4–8 years enrolled in the INMA cohort (female/male ratio= 0.87; n=109/125). Hg^R^ COPs were recovered from most faecal samples (185/232, 79.7%), with Hg^R^ Enterobacterales detected in 57.3% of individuals (133/232). These were frequently co-recovered with Hg^R^ enterococci and/or staphylococci (106/133, 79.7%), irrespective of host Hg burden (**Figure 1a-b**). Growth on gradient plates progressively declined along increasing Hg concentrations, revealing heterogeneous populations with variable Hg²⁺ susceptibilities (**Figure 1c-e**). Distinct Hg resistance phenotypes and genotypes frequently co-occurred within individual samples.

**Figure 1.**
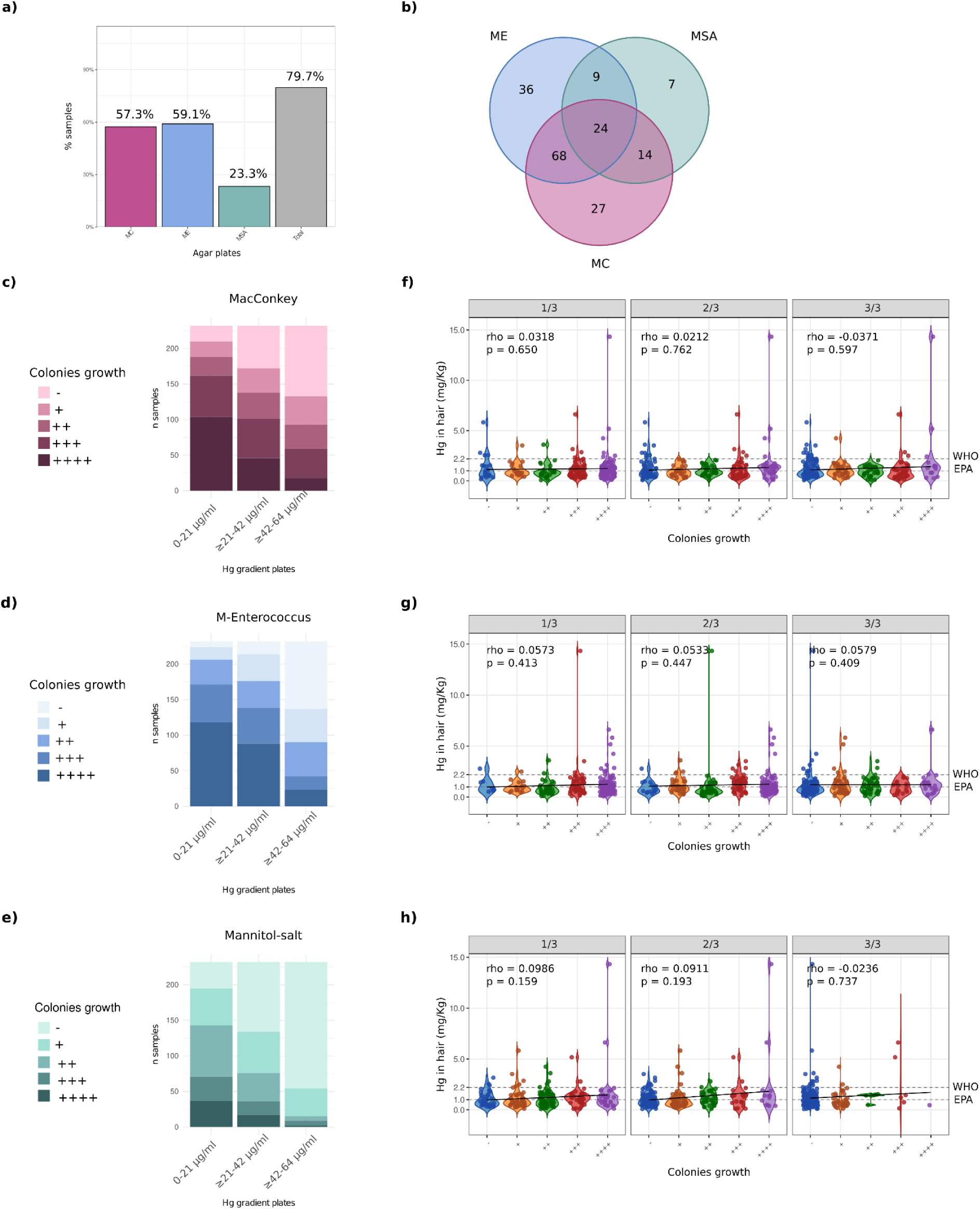
Faecal carriage of mercury resistance in children with exposure to pollutants. Samples containing Hg^R^ isolates from distinct taxonomic groups were identified based on growth on selective media: MacConkey (MC), m-Enterococcus (ME) and mannitol salt agar (MSA). **(a)** Percentage of samples yielding Hg^R^ Enterobacterales, enterococci, and staphylococci recovered on MC, ME, and MSA, respectively. The three categories are not mutually exclusive, as individual samples may be positive on more than one medium. Each bar represents all samples positive on the corresponding medium, including those also positive on one or both of the other media. “Total” represents the percentage of samples positive on at least one of the three media. Because the categories overlap, “Total” is not the sum of the individual percentages and may exceed the percentage for any individual medium. **(b)** Number of samples (n = 185) containing Hg^R^ isolates on each selective medium (MC, ME, or MSA). **(c-e)** Distributions of bacterial growth on gradient-selective plates, illustrated as stacked bar charts and categorised as - (0 colonies), + (1-10), ++ (10-100), +++ (100-1000), and ++++ (>1000). When manual counting was not feasible because overcrowding and colony confluence exceeded the standard limit of 300 CFUs per plate, we used either a “grid-based estimation method” or the “drop plate method” to accurately quantify high bacterial densities ^30,31^. **(f-h)** The correlation between bacterial growth on gradient-selective plates and hair Hg concentration (mg·kg-1) is shown using violin plots. Horizontal reference lines indicate established Hg exposure thresholds defined by the EPA (1 mg·kg-1) and WHO (2.2 mg·kg-1). Linear regression lines are included to visualise the overall trend. Spearman’s rho and p values are indicated in each panel.

Hair Hg concentrations ranged from 0.08 to 14.3 mg·kg^-1^ (P50 = 0.95; 95% CI 0.83–1.08). Overall, 45% of children (96/209) exceeded the US EPA reference level (1 mg·kg⁻¹), whereas 7.1% (15/209) exceeded the WHO and PTWI (Provisional Tolerable Weekly Intake) thresholds (2.0 and 2.2 mg·kg⁻¹, respectively). No association was observed between Hg^R^ bacterial carriage and hair Hg levels. Consistent with this, Hg^R^ isolates were recovered across the full spectrum of Hg exposure and dietary patterns represented in the cohort, despite previous associations between these variables and hair Hg concentrations ^24^ (**Figure 1f-h; Supplementary Figure 1**, **Supplementary Figure 2**).

### Diversity of Hg resistance across Enterobacteriaceae

Hg^R^ isolates recovered on MacConkey agar gradient plates (concentration range: 0 to 64 mcg/ml) and representing unique PFGE types and phenotypes (n=171/247; 69.2%) spanned diverse species and lineages (**Supplementary Data 1**). *Escherichia* predominated (62%; 106/171), largely driven by a phylogenetically diverse *E. coli* population encompassing all major phylogroups including *E. albertii*, *E. hermannii*, and *E. fergusonii*. Other frequently detected genera included *Klebsiella, Enterobacter* and *Citrobacter*, whereas *Kluyvera* and *Leclercia* were only occasionally isolated. PCR screening and hybridisation with *mer*-specific probes identified *mer* determinants in 73% (126/171) of the isolates recovered from the upper plate zone with the highest Hg concentrations. Of these, 79 were PCR-positive and 47 were detected by hybridisation. In contrast, 27% of phenotypically Hg^R^ isolates were negative in both assays.

PFGE of XbaI-digested genomic DNA revealed extensive genomic diversity. Whole-genome sequencing of a sample of 88 representative isolates further revealed a broad phylogenetic diversity among the Hg^R^ isolates (**Figure 2**). The complete *mer* operon was identified in 71 isolates, while 17 isolates carried partial or no detectable *mer* genes under the conditions tested.

**Figure 2.**
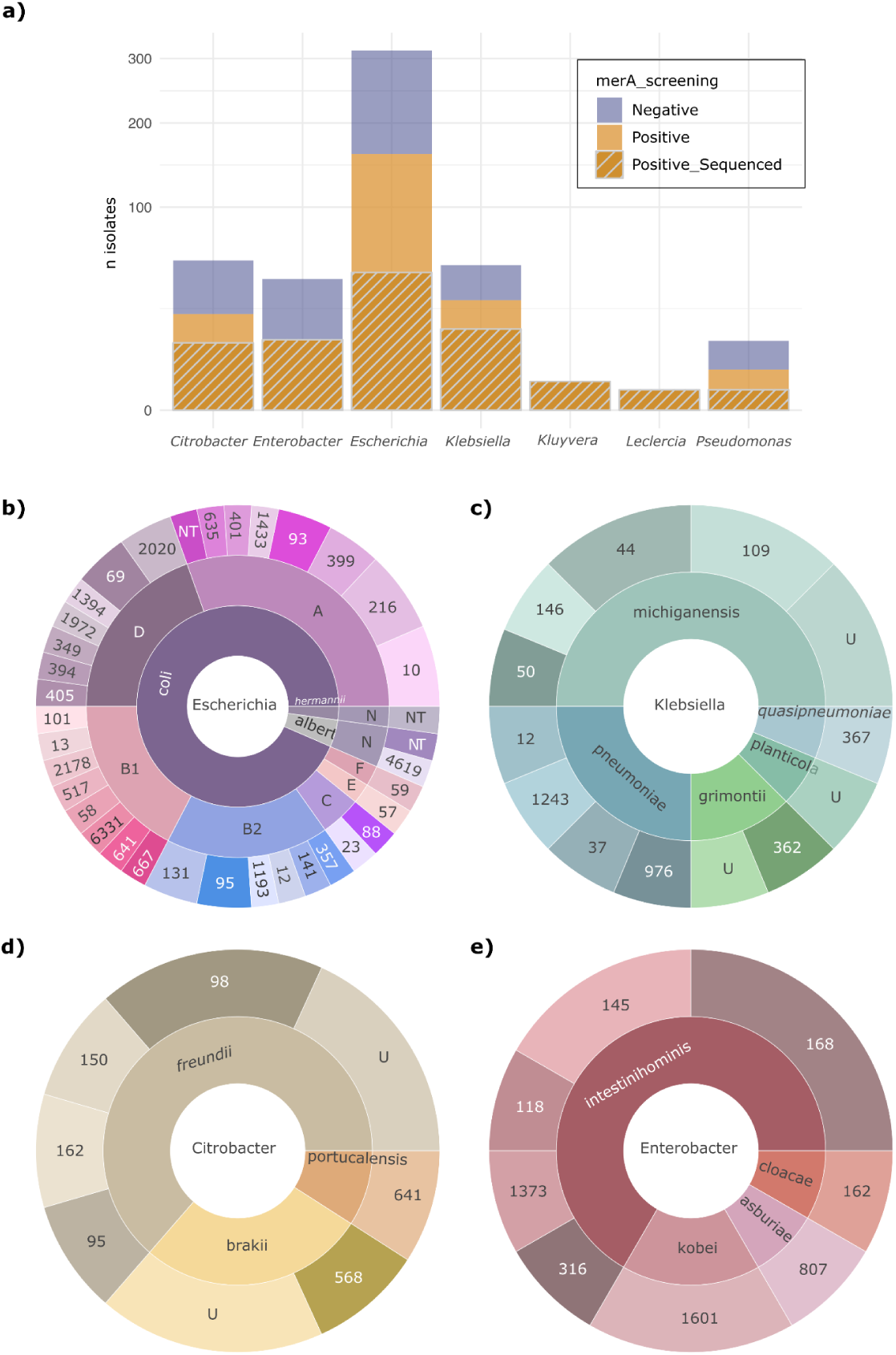
Taxonomic diversity of faecal mercury-resistant Enterobacterales. **(a)** Taxonomic distribution of *mer*A-positive Enterobacterales isolates recovered from gradient plates (n = 175), as determined by PCR or hybridisation. The y-axis shows the number of samples using a *square-root scale* to compress very large values and expand smaller ones. This transformation makes it easier to visualise and compare small numbers without large data points distorting the chart. **(b-e)** Relative frequencies of species and sequence types (STs) among whole-genome-sequenced isolates across different genera, shown as sunburst charts. The inner ring represents the genus, the second ring the STs for all genera except *Escherichia*, in which it represents the phylogroup, and the third ring shows the ST within each phylogroup. “U” indicates untyped STs. Among all unique isolates **(a)**, *Escherichia* predominated (106/171, 62%), largely driven by a phylogenetically diverse *E. coli* population (n = 101; phylogroups A, n = 36; B1, n = 23; B2, n = 18; C, n = 3; D, n = 15; E, n = 2; F, n = 3; untypeable, n = 1). Additional *Escherichia* species included *E. albertii* (n = 3), *E. hermannii* (n = 1) and *E. fergusonii* (n = 1). Other genera comprised *Klebsiella* (n = 21), *Enterobacter* (n = 21), *Citrobacter* (n = 20), *Pseudomonas* (n = 3), *Kluyvera* (n = 2) and *Leclercia* (n = 1). Whole-genome sequencing of 88 representative isolates included 43 *E. coli* (phylogroups A, n = 14; B1, n = 8; B2, n = 8; C, n = 2; D, n = 9; E, n = 1; F, n = 1), two *E. albertii*, one *E. hermannii*, 16 *Klebsiella spp*., 12 *Enterobacter spp*., 11 *Citrobacter spp*., two *Kluyvera spp.* and one *Leclercia adecarboxylata*. Additional strain data are shown in **Supplementary Data 1**.

### Diversity of Hg resistance determinants and *mer* operons

A total of 79 *mer* operons were identified across 71 isolates, including seven isolates carrying two distinct operons. They were grouped into three major backbones, namely, *merRTPCADE, merRTPFADE,* and *merRTPAGBD* (**Figure 3**). All carried *merA*, encoding a Hg reductase, and were therefore predicted to detoxify inorganic Hg by reducing toxic Hg²⁺ to volatile Hg⁰ (71/88 sequenced isolates, 80.7%). Four *mer* operons additionally carried *merB,* encoding an organomercurial lyase that cleaves the carbon–mercury bond, and three carried *merG,* encoding a periplasmic protein that reduces cell permeability to organomercurials, indicating an expanded capacity for organomercurial resistance and detoxification ^17^.

**Figure 3.**
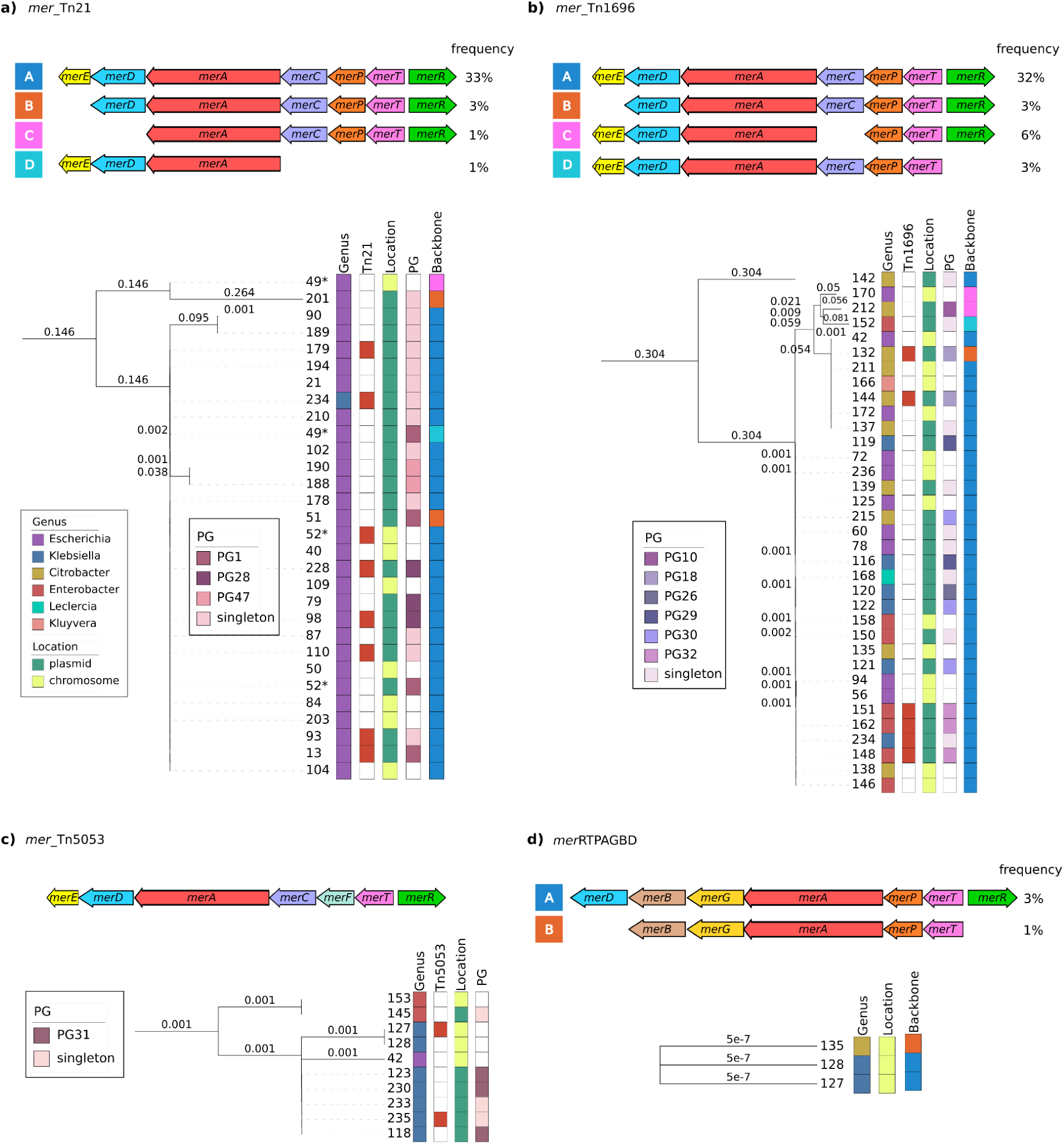
Diversity of operons conferring acquired mercury resistance. Schematic representations of the Mer operon architectures are shown and classified into four subgroups according to gene composition and amino acid sequence similarity to widespread **Hg^R^** transposons: *mer*_Tn*21*_ **(a)**, *mer*_Tn*1696*_ **(b)**, *mer*_Tn*5053*_ **(c)**, and Mer_RTPAGBD_ **(d)**. Operons were assigned to the same subgroup when sharing at least 85% amino acid sequence similarity, at both individual protein and operon levels. Although the MerRTPAGBD backbone showed protein similarity with the *mer*_Tn*5053*_ subgroup, it was treated as a distinct operon because its amino acid sequence similarity was <85%. Operon sequences in panels a, c, and d were highly similar, whereas those in panel b split into two subgroups. Arrows indicate gene content and orientation, and percentages denote the frequency of each configuration. The accompanying phylogenetic tree highlights sequence diversity across isolates; asterisks indicate sequences derived from the same isolate. The phylogenetic tree was plotted and annotated using iTOL. Seven isolates carried multiple *mer* operons, either on different plasmids or in both plasmid and chromosomal locations: p*mer*_Tn*21*_ + Chr*mer*_Tn*21*_ (2 *E. coli*); p1*mer*_Tn*21*_ + p2*mer*_Tn*1696*_ (1 *Klebsiella*); Chr*mer*_Tn*5053*_ + ChrMerRTPAGBD (2 *Klebsiella*); Chr*mer*_Tn*1696*_ + Ch*rmer*_RTPAGBD_ + p*mer*_RBDE_ (1 *Citrobacter*); Chr*mer*_Tn*1696*_ + Chr*mer*_Tn*5053*_ (1 *E. coli*).

The *merRTPCADE* backbone predominated (62/88 isolates; 70.5%) and comprised two subgroups, *mer*_Tn*21*_ (28 isolates) and *mer*_Tn*1696*_ (35 isolates), defined based on >85% amino acid sequence similarity of individual Mer proteins and the corresponding genetic organisation of characterised *mer* operons from Enterobacterales Hg-Tns.

Most *mer_T_*_n*1696*_ operons were highly conserved (25/35 isolates), although seven shared a distinct variant that differed in the sequences of three proteins (<85% amino acid sequence similarity). The remaining three corresponded to a Tn*1696*–Tn*5053* hybrid, *mer*_Tn*6346*_, and *mer*_Tn*6203*_. The *merRTPFADE* backbone (n = 10) was exclusively associated with *mer*_Tn*5053*_. The third backbone, *merRTPAGBD* (n = 3), represented a previously undescribed organomercurial-degrading operon. The distribution of *mer* operons differed among genera. While *mer*_Tn*21*_ was largely restricted to *Escherichia*, *mer*_Tn*1696*_ was detected across all genera, and *mer*_Tn*5053*_ and MerRTPAGBD backbone were primarily detected in *Klebsiella*.

Operons within each subgroup showed variable sequence conservation and indel patterns, and were located on both plasmids (p*mer*) and/or chromosome (Chr*mer*), although plasmid-borne operons predominated (63.3% versus 36.7%). Complete Hg transposons were identified in a subset of isolates (**Figure3**).

### Metal resistome of Hg^R^ Enterobacteriaceae

Beyond *mer* operons, Hg^R^ isolates carried a broad repertoire of acquired (a) and intrinsic (i) MRGs targeting copper, silver, nickel, iron, and arsenic. The genetic organization and functions of these metal resistance systems has been extensively described in previous publications ^4,5,17,32^ The distribution of these MRGs showed strong species-level structuring (**Supplementary Figure 3 and Supplementary Data 2)**. Non–*E. coli* isolates were enriched in *pco* and *sil* systems, whereas *E. coli* strains formed two major clusters, one enriched in siderophores (aerobactin, salmochelin, yersiniabactin) and iron uptake systems (*sitABCD*, *fecIRABCDE*), and a second characterised by the presence of *pco* and *sil* operons but lacking these iron-scavenging determinants.

Copper and silver resistance genes (*pcoABCDRSE* and *sil*PABFCRSE) were present in 52% of isolates (46/88), where they consistently co-occurred. They exhibited greater sequence diversity than *mer* operons, including multiple indel-based variants (**Supplementary Figure 4).** In *Klebsiella,* both operons were plasmid-borne, whereas in *Escherichia*, *Enterobacter*, and *Citrobacter,* were located on either plasmids or chromosomes. Ten isolates (five *E. coli* and five *Citrobacter*), carried chromosomal regions containing *pco*, *sil,* and *cus* genes, consistent with the Copper Homeostasis and Silver Resistance Island (CHASRI) ^33^.

Iron-acquisition determinants were widespread but unevenly distributed. The ferrous iron transporter *sitABCD*, present in 74% of isolates, was detected across all genera, and occurred on plasmids in *Escherichia* despite being typically chromosomal in Enterobacteriaceae. The ferric citrate uptake system *fecIRABCDE* was identified in 61% of isolates and frequently co-occurred with *sitABCD*. Whereas the *fec* system was exclusively chromosomal in *Escherichia* and *Citrobacter*, it was located on either plasmids or chromosomes in *Klebsiella* and *Enterobacter*, indicating genomic mobility in these genera. Siderophore systems showed strong taxonomic specificity. Aerobactin (*iucABCD-iutA*) and salmochelin (*iroBCDEN*) were restricted to *E. coli*, whereas yersiniabactin (*ybt*PQ) was present in both *E. coli* and *Klebsiella*. Arsenic-resistance genes were the most prevalent MRGs. Chromosomal *arsRBC* genes were detected in 88-97% of isolates, consistent with the widespread presence of the *arsRBC* operon. In contrast, *arsA*, *arsD* and *arsH* were less frequent (39-43%) and were located on both plasmids and chromosomes, consistent with the acquisition of additional *ars* operons (*arsRDABC*, *arsRDABCH*) (see below).

In addition, isolates carry a broad repertoire of iMRGs involved in metal homeostasis and stress responses that are often misclassified as canonical antimicrobial resistance genes ^4^ (**Supplementary Figure 3**). Conserved chromosomal systems mediating the uptake and trafficking of essential divalent cations, including *zupT*, *mnt* and *nik* transporters, were common across the collection, although *nik* genes showed marked taxonomic structuring and were largely restricted to *Escherichia* and *Klebsiella*. Core copper-homeostasis determinants (*cue*, *cut*) were detected in most isolates, whereas the Cu/Ag efflux system *cus* exhibited a narrower distribution and was nearly ubiquitous only in *E. coli*. Genes involved in arsenic uptake and detoxification, including phosphate transporters and *glpF*, were broadly conserved. Global stress-response genes associated with metal tolerance, such as *dsbAB* and *robA*, were also widespread.

### Antibiotic resistome of Hg^R^ Enterobacteriaceae

Most Hg^R^ Enterobacteriaceae isolates were resistant to at least one antibiotic (72%; 123/171), although resistance profiles varied markedly among genera. *Escherichia* showed the broadest resistance profiles, encompassing β-lactams, aminoglycosides, quinolones, trimethoprim, tetracycline, and sulfonamides, whereas *Enterobacter* and *Citrobacter* were primarily resistant to β-lactams, and *Klebsiella* showed limited resistance, largely restricted to ampicillin (**Supplementary Data 3**).

Acquired resistance to multiple antibiotic classes (β-lactams, aminoglycosides, trimethoprim, sulfonamides, tetracyclines, colistin) was associated with aARGs located on widespread transposable units present in both pristine and anthropogenically impacted environments ^32^. They include class 1 integrons (*aadA, dfrA, sul1*, *bla*_OXA-1_), *sul2*, *tetA*/Tn*1721*, *bla*_TEM_/Tn*3, aph*(*3′′*)-*Ib-aph*(*6′*)-*Id* (also called *strA-strB*). Four isolates carried the contemporary aARG *mcr*-9.1, a plasmid-mediated colistin-resistance determinant frequently associated with MRGs and silently disseminated in recent years across OneHealth sectors on antibiotic resistance plasmids ^34^. In contrast, β-lactam resistance was explained by species-specific chromosomal iARGs, including *bla*_SHV_ and *bla*_OXY_ in *Klebsiella*, *bla*_CMY_ in *Citrobacter*, *bla*_ACT_ in *Enterobacter*, *bla*_AMPC_ in *Escherichia*, and *bla*_CTX-M_ in *Kluyvera* (**Supplementary Figure 5; Supplementary Data 3**). Overall, the antibiotic resistome showed strong genus-level structuring and was dominated by iARGs and widely disseminated aARGs ^32^ rather than by recently emerged clinically relevant resistance determinants encoding first-line antibiotics such as third generation cephalosporins or carbapenems.

### The plasmidome of Hg^R^ Enterobacteriaceae

Plasmids in our dataset were characterised using three complementary and widely used plasmid-typing approaches, COPLA^35^, replicon-typing ^36^, and network-based plasmid clustering (see Materials and Methods). The latter enables reference-independent classification of plasmid sequences and is particularly useful for previously uncharacterised or hybrid plasmids that are difficult to categorise using reference-based approaches, as observed here and in other large plasmid collections ^37,38^.

Network-based plasmid clustering identified a plasmidome of 363 plasmids across 88 HgR genomes analysed, of which 163 clustered into 52 plasmid groups (PGs), while 200 remained as singletons. aMRGs were detected in 52 plasmids (10 PGs and 30 singletons), aARGs in 28 (4 PGs and 17 singletons), and both aMRGs and ARGs in 35 (4 PGs and 16 singletons) (**Figure 4, Supplementary Data 4**).

**Figure 4.**
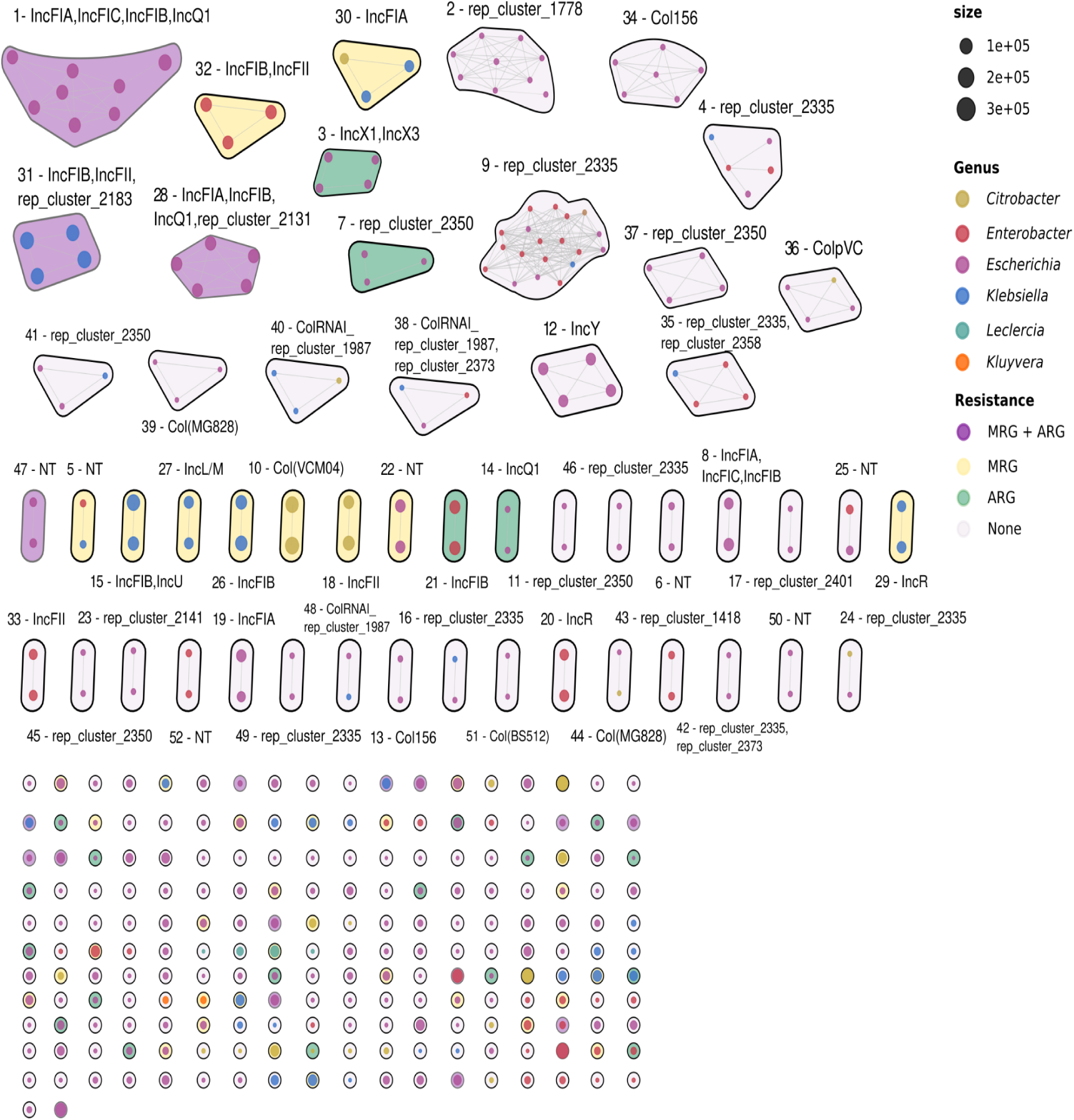
The plasmidome of HgR Enterobacterales. Each node represents a plasmid, coloured by genus, and node size is proportional to plasmid length. Plasmid groups (PG) are clusters containing at least two members, whereas ungrouped plasmids are considered “singletons”. PGs and singletons were assigned arbitrary numbers, and their replicon content is indicated in the labels. Parentheses indicate variable replicon types within the group, and asterisks denote PGs in which the replicon type may vary among plasmids. Node colours denote the resistance profile of each PG or singleton: MRGs, ARGs, both, or no resistance genes. The complete plasmidome dataset identified across the genomes analysed is provided in **Supplementary Data 4**.

Larger plasmids more frequently carried MRGs, either alone or in combination with ARGs. Plasmids carrying MRGs and ARGs were most common in *Escherichia* and *Klebsiella*, whereas ARG-only plasmids were not detected in *Klebsiella* or *Citrobacter*. Across genera, isolates carried between one and more than seven plasmids, with *Escherichia* and *Enterobacter* showing the highest mean plasmid counts per isolate (**Figure 5**).

**Figure 5.**
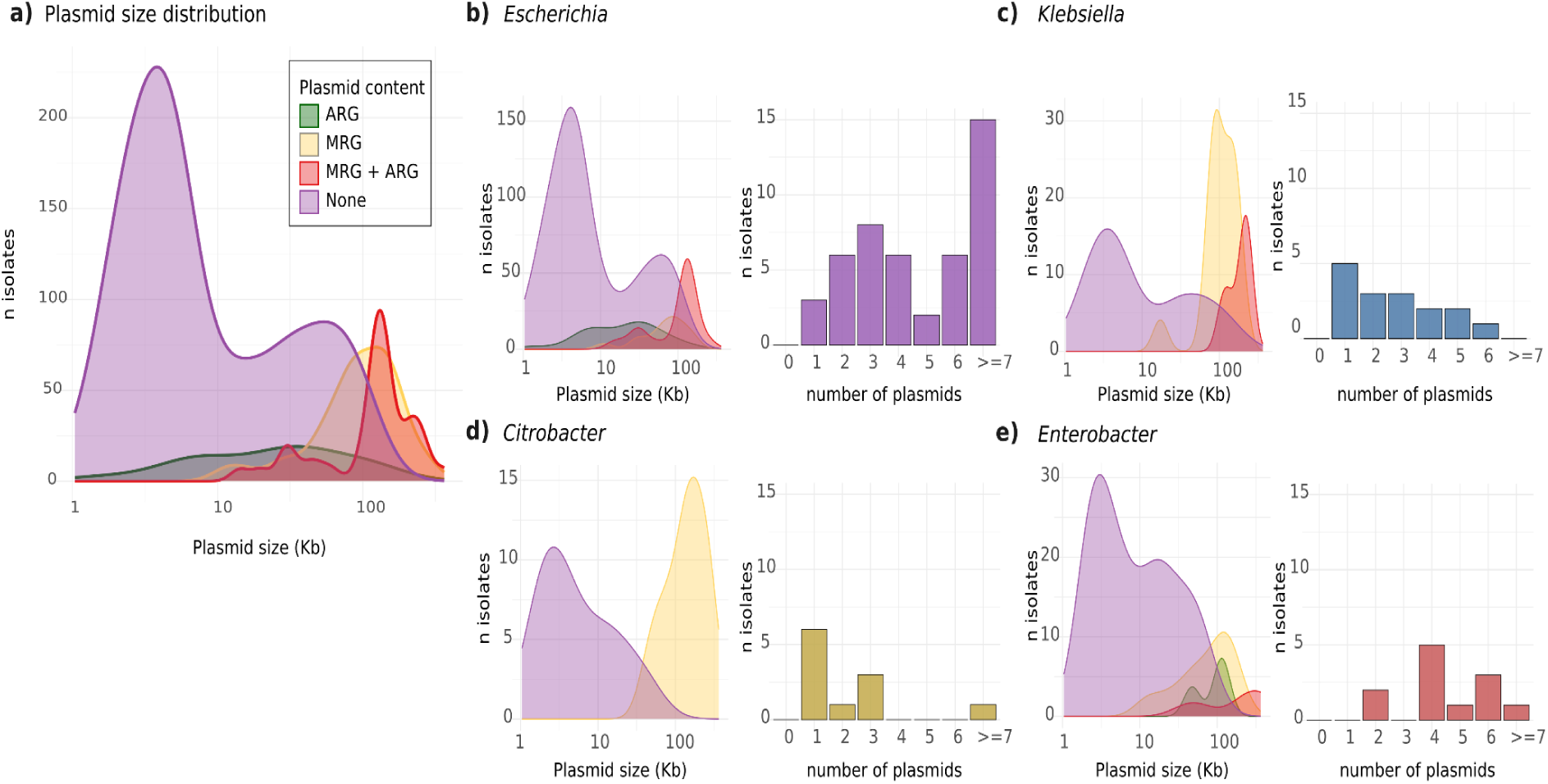
Size and number distributions of plasmids in HgR Enterobacterales. **(a)** Estimated plasmid size distribution on a log10 scale. Panels **b-e** show, for each genus, a kernel density distribution of plasmid sizes and a histogram of the number of plasmids per isolate: *Escherichia* **(b)**, *Klebsiella* **(c)**, *Citrobacter* **(d)** and *Enterobacter* **(e).** In the kernel density plots, colour indicates the presence of metal resistance genes (MRG), antibiotic resistance genes (ARG), both (MRG + ARG), or neither. Plasmid data are provided in **Supplementary Table 4**.

Hg resistance was predominantly plasmid-borne (62.8%), distributed across 10 PGs and 26 singletons, with marked taxonomic structuring (**Figure 6, Supplementary Data 5 and 6**). Hg^R^ plasmids were enriched in F plasmids, corresponding mainly to COPLA flagship groups characteristic of *E. coli* (PTU-E81 and PTU-FE) and *Klebsiella* (PTU-FK). Other incompatibility groups, including K, I1, HI2, Q, and R, were also detected and corresponded to more “generalist” PTU groups such as PTU-B/O/K/Z (PG82), PTU-I1 (PG85) and PTU-HI2 (PG73). Plasmids from non-*E. coli* isolates were frequently unassigned by reference-based typing, while chimeric plasmids sometimes received incongruent PTU assignments. For example, PG29 is an 88kb hybrid IncFIA-IncR *Klebsiella* plasmid assigned by COPLA to either PTU-E71, typically associated with small Col4401 plasmids, and to cPTU-6635_3 (**Supplementary Data 4 and 5)**.

**Figure 6.**
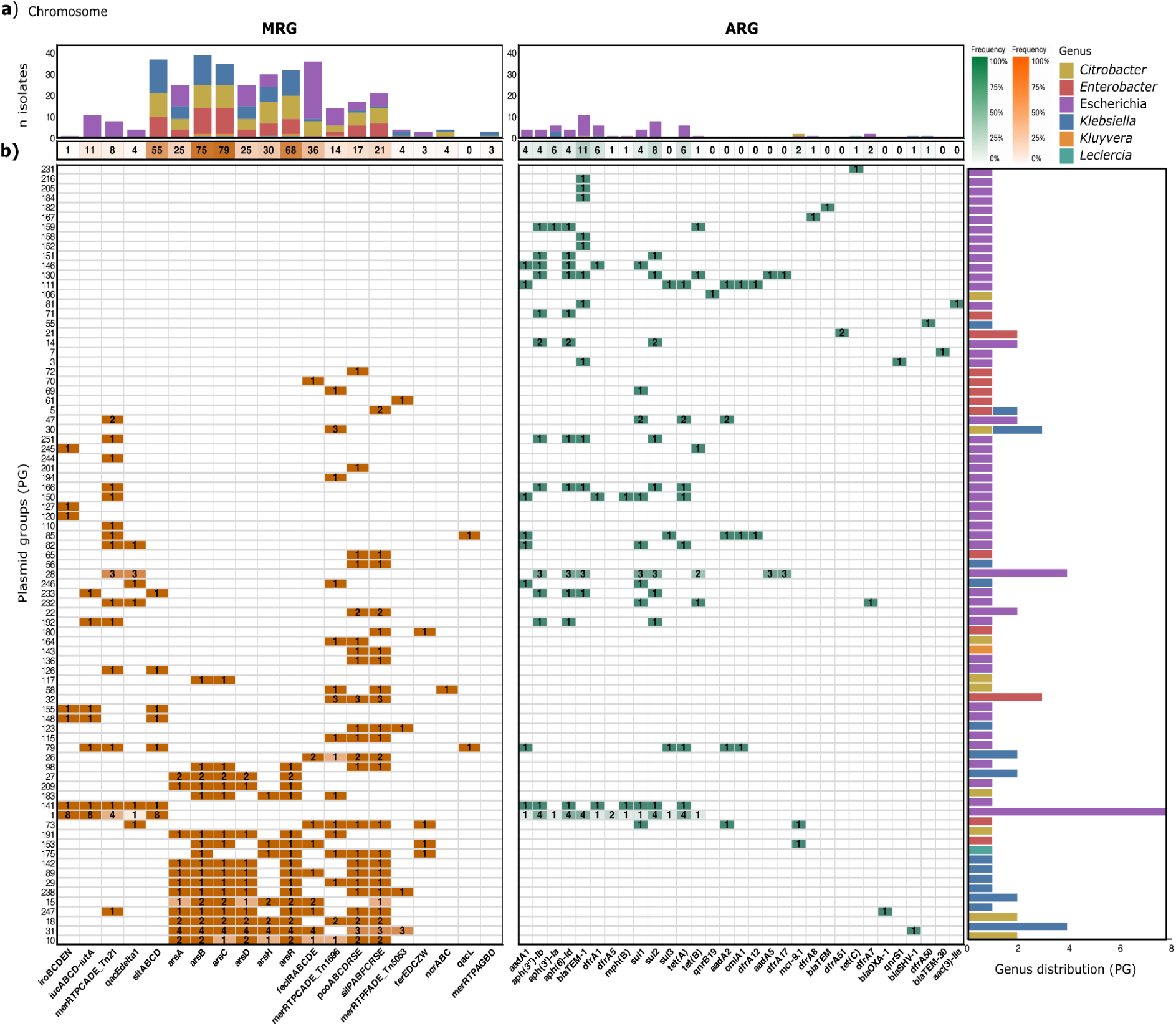
Genomic location of acquired genes conferring resistance to metals and antibiotics. The figure shows the number of isolates carrying acquired metal resistance genes (aMRGs) and acquired antibiotic resistance genes (aARG) on the chromosome (Panel a) and across plasmid groups (PGs) (Panel b). The marginal bar charts show the number of isolates from each species that contain each feature. MRGs and ARGs data are provided in **Supplementary Data 2 and 3**, respectively.

The *E. coli-*associated *mer*_Tn*21*_ operon was mainly carried by F-type plasmids belonging to PTU-E81 or PTU-FE macrolineages and also by IncI1gamma (PTU-I1) or IncK (PTU-B/O/K/Z) plasmids, which have been associated with the food chain ^39^. Several F plasmids (PG1, PG79, PG141, PG192) also consistently carried additional adaptive traits, including iron acquisition systems such as aerobactin, salmochelin, and Fe²⁺/Mn²⁺ transport loci. All except PG192 belonged to the PTU-E81, a lineage previously reported to evade restriction-modification systems and occur across diverse *E. coli* hosts ^40^. PTU-FE plasmids (PG28, and the singleton PG232) additionally carried ARGs. In other *Escherichia* species, the *mer*_Tn*21*_ operon was located on plasmids lacking reference-based assignments.

In non-*E.coli* species, the *mer*_Tn*1696*_ and *mer*_Tn*5053*_ operons were also mainly associated with F-type plasmids. Most lacked reference-based assignments in plasmid databases and carried multiple metal-resistance systems in addition to *mer* **(Supplementary Data 5 and 6)**. Some also carried additional adaptive traits, including *hsp20*, *clpK*, and the heat-resistance locus (LHR), as observed in PG31, which belongs to the PTU-FK macrolineage, and PG32, which received an incongruent PTU by COPLA (**Supplementary Figures 8 and 14**).

Despite this diversity, Hg^R^ plasmids recovered from children in different regions often showed strikingly high sequence conservation, with up to 99% coverage and 100% nucleotide identity, and frequently corresponded to established plasmid lineages. These included the globally distributed *E. coli* IncF–IncQ1 hybrids PG1 (PTU-E81) and PG28 (PTU-FE), previously reported in clinical high-risk *E. coli* lineages ^41,42^ (**Supplementary Figures 6 and 7**).

Other highly conserved plasmids carried multiple MRGs, including PG18 and PG29 (*mer*+*pco+sil+ars*), PG26 and PG10 (*mer+pco+sil+fec*), and PG31 (*mer*+*pco+sil+fec+ars*). PG31 belonged to the PTU-FK macrolineage, whereas the others either lacked any reference-based assignments or received incongruently COPLA classifications (**Supplementary Figures 8-14**) ^43^. Additional examples included IncK (PG82), *sul3*-IncIγ (PG85), and *mcr 9.1*-IncHI2 (PG73) plasmids, all previously reported in food-associated environments ^34,39^. Notably, both *mer*-positive and *mer*-negative variants occurred within otherwise highly conserved plasmid backbones, both in our collection and in public databases (**Supplementary Figures 6-8, 10-12, 14; Supplementary Data 4-6).**

### Correlation analysis of MRGs

Co-occurrence analysis (**Figure 7)** showed that *mer* operons occurred either as the sole MRG system, predominantly in *E. coli,* or in combination with additional MRGs, particularly in non-*E. coli* species. These combinations were typically associated with widely disseminated MGEs, most often plasmids carrying additional adaptive traits.

**Figure 7.**
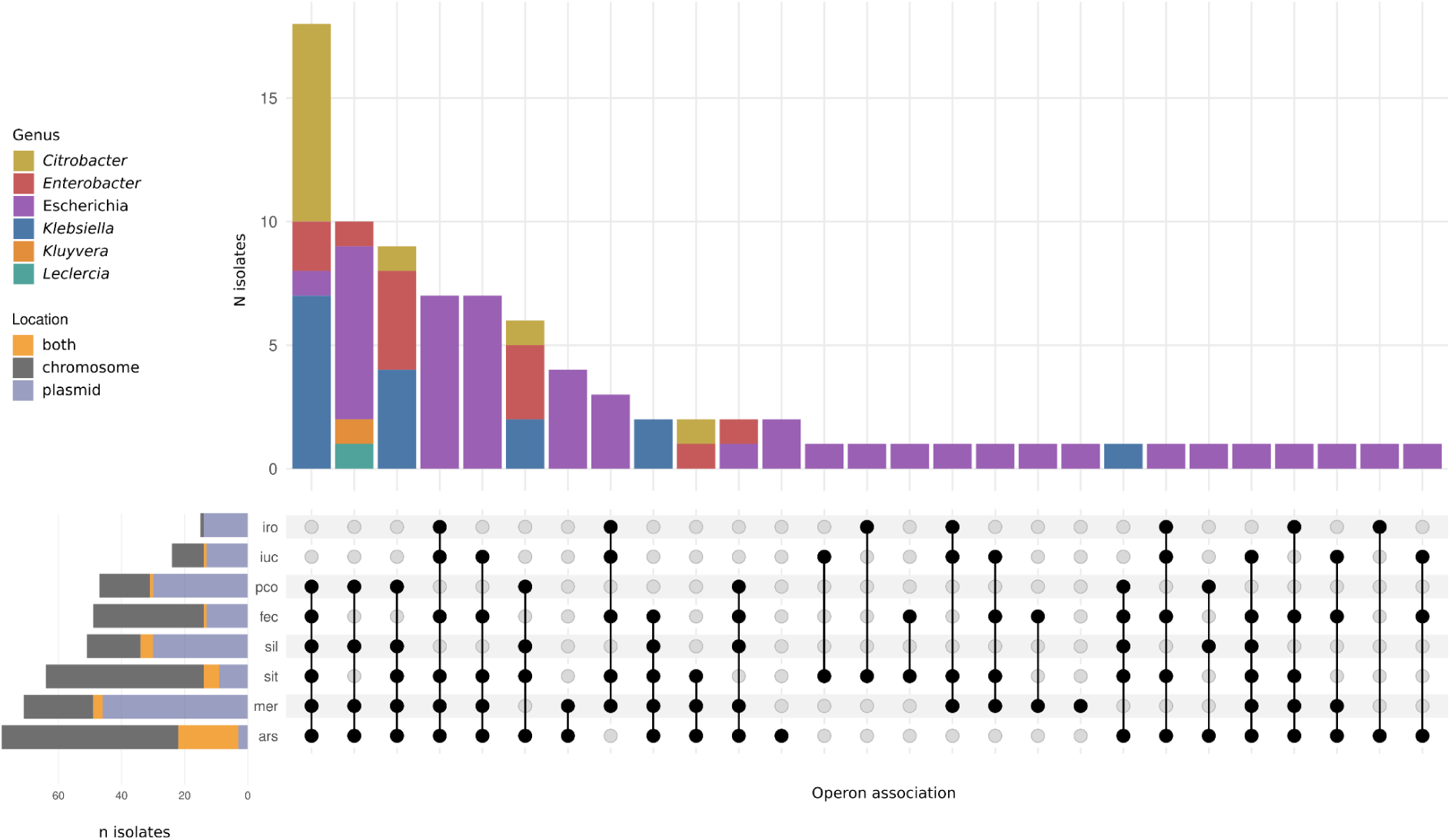
Distribution of metal resistance gene (MRG) profiles across bacterial genera. The upper bar chart shows the number of isolates from each genus carrying individual or combined MRG systems. The marginal bar charts on the left indicate the genomic location of each determinant and the number of isolates from each species carrying the corresponding feature (Hg: *mer*; As: *ars*; Cu: *pco*, *sil*; Fe: *fec*, *iro*, *iuc*, *sit*). Metal systems were represented by their consensus acronyms (*mer*; *ars, pco*, *sil, fec*, *iro*, *iuc*, *sit*), with each metal acronym encompassing the diversity of corresponding systems described in the previous section. Additional data are provided in **Supplementary Table 7**.

Resistance determinants were frequently co-located on the same genetic element or distributed across multiple plasmids and chromosomes within the same isolate (**Supplementary Table 7**), suggesting repeated independent acquisition events on available units across the gut microbiota. Isolates carrying more than three or four MRG systems were uncommon. Co-resistance between ARGs and MRGs was frequent and occurred either within the same genetic element or across distinct plasmid and chromosomal loci (**Figure 6**).

### *Mer* operon occurrence in public databases

Comparison with the AllTheBacteria database (https://allthebacteria.org/) ^43,44^ showed that *merA* was more frequently detected in our Hg-selected collection than in the public genomes across the analysed genera (Fisher’s exact test, p < 0.05; **Supplementary Data 8**). The difference was particularly pronounced in *Escherichia* and *Klebsiella*, where *merA* was absent from the public genomes in several taxa. Although *pcoA*, *silA* and *arsA* were also more frequently detected in our collection, their differences were less pronounced. In contrast, siderophore-associated genes were predominant in *Escherichia* and absent in other genera (**Supplementary Figure 15**). This comparison should not be interpreted as a measure of resistance prevalence, given the Hg-selection of our collection and the heterogeneous origins and study-specific selection criteria of the public genomes. Rather, it highlights the limited representation of Hg-resistance determinants in public genomic datasets, where such determinants may not have been specifically targeted or characterised, pointing to a potentially underrecognised or “hidden” Hg-resistance resistome.

## Discussion

This study provides the most comprehensive characterization of functional Hg resistance in the human gut microbiota to date. Using a Mediterranean pediatric cohort as a model, we show that highly conserved *mer* operons circulate across taxonomically structured and clonally diverse bacterial communities through a broad range of MGEs, including both rare and widely distributed plasmid lineages. These findings reveal the persistence and dissemination of Hg resistance determinants within human-associated microbial communities and highlight their potential relevance to the broader ecology of AMR.

A first indication of the ecological success of Hg resistance was the widespread carriage of Hg^R^ bacteria across the INMA cohort. Similar frequencies, whether estimates from *E. coli* or from phenotypically Hg^R^ COP species as a whole, have been reported in geographically and socioeconomically distinct cohorts, including Amazonian indigenous communities and urban populations ^32,45^. Historically, high prevalence of Hg^R^ bacteria has been attributed to direct or indirect Hg exposure from sources such as artisanal gold mining, fish consumption, dental amalgams, antiseptics, or pharmaceuticals ^15,16,28,45^. However, associations between Hg exposure and bacterial Hg resistance have been inconsistent across studies. Despite substantial differences in host populations, ecological settings, exposure levels, and approaches used to quantify *mer* genes, Hg resistance consistently emerges as a pervasive feature of human-associated microbial communities. Together, these observations suggest that contemporary Hg resistance reflects not only current exposure but also the persistence and circulation of resistance determinants within interconnected microbial communities.

The INMA cohort provides a relevant setting in which to examine this relationship, as elevated Hg exposure has previously been reported in pregnant women and in their offspring, who retain measurable body burdens throughout childhood despite a gradual decline with age ^24–27^. Nevertheless, Hg^R^ bacterial carriage was not associated with either hair Hg concentrations or fish consumption in our cohort, which further supports a decoupling between host-level exposure metrics and the ecological distribution of bacterial Hg resistance. Hg biomarkers often integrate multiple and overlapping sources of exposure and therefore capture only a fraction of the variation in individual exposure ^14^. Moreover gut microorganisms can influence Hg speciation, absorption, and excretion, potentially further decoupling host Hg burden from microbial responses ^46,47^. Together, these findings suggest that ecological and evolutionary processes may contribute substantially to the persistence of Hg resistance beyond contemporary exposure. The strong taxonomic structuring observed here further supports an ecological component, with *Escherichia* lineages frequently associated with iron-acquisition systems and siderophores, whereas non-*Escherichia* taxa more often carried multimetal-resistance determinants, including *pco*, *sil* and *ars* operons. These contrasting profiles may reflect differences in bacterial ecological niche and host adaptation, contributing to the uneven distribution of resistance determinants across gut-associated bacterial populations.

The concurrent circulation of a diversity of *mer* operons across a broad spectrum of both rare and widely distributed plasmids resembles the long-tail abundance patterns observed in ecological communities, in which numerous individually rare entities coexist with a limited number of highly successful lineages ^48^. This pattern suggests that the contemporary landscape of *mer* operons reflects the combined effect of historical dispersal, ecological filtering and long-term persistence of successful MGEs, rather than recent selection alone. Consistent with this view, *mer* operons would have been independently acquired by diverse plasmid families, with a distribution across species shaped by genetic compatibility, host ecology and bacterial lifestyle, as also observed for ARGs ^49^. Ecological coexistence theory provides a useful framework for interpreting this pattern, as HGT can maintain adaptive functions across multiple taxa and partially decouple community functions from taxonomic composition over time ^50^. Such functional redundancy may promote the persistence of adaptive functions even under antimicrobial selection by allowing multiple taxa to retain or acquire functions favoured by the prevailing environment ^51^. Although ecological models, also applied to antibiotic resistance, have proposed that increasing diversity can constrain the resistance spread through dilution effects, frequent HGT and environmental fluctuations may weaken this constraint by redistributing adaptive functions among taxa ^51,52^. Metal resistance provides a particularly relevant context for evaluating these dynamics. Unlike the often short-term and episodic selective pressures imposed by antibiotic use, metal exposure is ancient and persistent but highly heterogeneous across space and time, and with indirect impacts on humans. Hg contamination also frequently co-occurs with other metals, including copper and silver, through shared environmental sources ^53^ and their long histories of industrial and medical use ^14,15^. Such conditions may favour the long-term maintenance and redistribution of conserved metal-resistance operons ^33,54^.

The *mer* operon architectures identified here, including those associated with Tn*21*, Tn*1696,* and Tn*5053*, have persisted from the pre-antibiotic era to the present and are frequently linked to plasmids involved in the emergence and spread of antibiotic-resistant Enterobacteriaceae ^29,54–60^. Our results extend these observations by showing that diverse *mer* and other metal-resistance operons circulate within a single pediatric population across plasmids of multiple incompatibility groups. These include F-plasmid groups nested within species-associated macrolineages such as PTU-FE, PTU-FK, primarily linked to antibiotic resistance ^61^, PTU-E81, associated with host colonisation and immune defence ^40,42^ as well as broadly distributed plasmid groups such as IncI1gamma (PTU-I1), IncK (PTU-B/O/K/Z), or IncHI2 (PTU-HI2), commonly detected in foodborne and animal-associated bacteria ^34,39^. We also identified widespread multimetal-resistance plasmids from non-*E. coli* species that could not be assigned, or were inconsistently classified by current plasmid-typing approaches, consistent with previous plasmidome studies ^37^. Together, these results highlight the ecological diversity of MGEs carrying Hg^R^ determinants in a single population and support the long-term persistence and diversification of successful MGE lineages^48,62^, consistent with recent plasmidome analyses showing continuity between contemporary plasmids and those predating the antibiotic-era ^38^. Rather than acting solely as transient vehicles of HGT, plasmids and transposons provide genetic scaffolds on which adaptive traits, including MRGs and ARGs, can be repeatedly acquired, reshuffled, and disseminated. Thus, the persistence of Hg resistance may reflect not only metal-mediated selection but also the ecological success and evolutionary history of the MGEs carrying these determinants.

The frequent co-occurrence of multiple MRGs may promote genetic linkage and co-selection, favouring their persistence across environmental reservoirs and dissemination through food webs into human-associated bacterial communities. This process may be further reinforced by the use of metals as feed additives and growth promoters in agrifood systems. Co-selection between MRGs and ARGs is particularly relevant in this context. Several widespread aARGs identified here, including *sul2*, *strA-strB*, and *aph(3′)-III*, are also prevalent in Western human gut microbiomes and have been associated with MGEs from ancient or minimally impacted environments^32^. Other aARG linked to the widespread integrons detected here, including *dfrA1-aadA1*, *dfrA17-aadA5*, and *dfrA21-aadA3* repeatedly co-occurred with aMRGs (*mer*, *pco*, *sil*) in our previous study of the resistome of Indigenous populations ^32^. These observations are again consistent with a resilient mobile resistome in which individually variable resistance can nevertheless be maintained at the through shared genetic contexts. Nonetheless, more than 30% of *mer* operons and other MRGs were chromosomally encoded, with their distribution varying across taxa. This taxonomic structuring is consistent with patterns reported for ARGs in *Escherichia* and *Klebsiella* ^49^, indicating that resistance dissemination is shaped not only by plasmid-mediated linkage and co-selection but also by chromosomal localization and host-associated genomic context, which may constrain HGT.

In summary, these findings reveal a highly connected yet ecologically structured network of MRGs within the human gut microbiota. They extend exposure-based models of antimicrobial resistance by showing that resistance determinants can circulate across ecological compartments beyond contemporary exposure, highlighting the human gut microbiota as a reservoir connecting environmental and clinical resistance evolution. The occurrence of both *mer*-positive and *mer*-negative variants within highly conserved plasmid backbones further indicates repeated, independent acquisition of *mer* operons by successful plasmid lineages, underscoring the capacity of persistent MGE scaffolds to repeatedly recruit and redistribute adaptive resistance traits. In this context of extensive plasmid diversity and conservation, the complementary use of current plasmid classification approaches is essential to capture the full diversity of plasmid-mediated resistance and provide a more comprehensive view of the antimicrobial resistance landscape.

## Methods

### Study design and sampling

This study analysed 234 faecal samples collected between 2012 and 2013 from a cohort of children (4-8y, female-to-male, ratio= 0.87, 109/125). Participants were enrolled in the INMA project (<u>IN</u>fancia y <u>M</u>edio <u>A</u>mbiente, Childhood and Environment, https://www.proyectoinma.org/en/inma-project/general-overview/), a Spanish birth cohort study assessing the impacts of environmental pollutants in air, water, and diet during pregnancy and early childhood, with follow-up in young adulthood. Participants were children who lived in three geographically distant areas in Spain, namely Sabadell (161/234; 68.8%), Gipuzkoa (35/234; 15%), and Valencia (38/234; 16.2%). Variables used to assess the impact of Hg exposure on the carriage of Hg^R^ bacteria included Hg concentrations in hair, as well as dietary factors. Correlations between these variables and the growth of Hg^R^ isolates on Hg gradient plates were assessed using robust Spearman’s rank correlation.

Of the 234 collected samples, 232 were available for microbiological screening. Initial screening for Hg^R^ bacteria used gradient plates of MacConkey agar, m-Enterococcus, and Mannitol salt agar (MSA) (Becton, Dickinson and Company, Sparks, Le Pont de Claix, France) supplemented with HgCl₂ at concentrations ranging from 0 to 64 µg/mL ^63^. Colonies were initially counted manually. When manual counting was not feasible due to overcrowding and colony confluence, resulting in bacterial densities exceeding the standard upper limit of 300 CFUs per plate, colony numbers were estimated using either a grid-based estimation method or the drop plate method to ensure accurate quantification of high bacterial densities ^30,31^. Colonies growing in the upper zones of the gradient plates were subsequently plated on selective plates supplemented with 128 µg/mL HgCl₂ to confirm resistance. Hg^R^ colonies representing different morphotypes (based on size, colour, and shape) were identified by MALDI-TOF MS (MALDI Biotyper, Bruker, Billerica, MA, USA), using a reliability score >2 for species-level identification. Confirmed isolates were subcultured, and bacterial stocks were stored at −80 °C in 1 500 µL of Luria–Bertani broth supplemented with 15% glycerol for further analyses.

### Identification and preliminary characterisation of isolates

All Hg^R^ isolates were initially analysed by pulsed-field gel electrophoresis (PFGE) following PulseNet protocols (https://pulsenetinternational.org/protocols/pfge/). *E. coli* were assigned to major phylogenetic groups (A, B1, B2, C, D, E, and F) and B2 subgroups (B2-I to B2-X) using multiplex PCR schemes ^64,65^ and Hg^R^ isolates were screened for the *merA* gene and Hg transposons by PCR and Southern blot hybridisation ^58^. Isolates with distinct PFGE profiles were selected for whole-genome sequencing (WGS).

### Antimicrobial susceptibility

Susceptibility to 16 antibiotics was assessed by disk diffusion on Mueller-Hinton agar (Becton, Dickinson and Company, Sparks, Le Pont de Claix, France) using antibiotic discs (BioRad, Marnes-la-Coquette, France) at concentrations established by EUCAST guidance documents ^66^. The antibiotics tested were ceftazidime (CAZ), amoxicillin-clavulanate (AMC), cefotaxime (CTX), ampicillin (AMP), cefoxitin (FOX), meropenem (MEM), ciprofloxacin (CIP), nalidixic acid (NAL), gentamicin (GEN), kanamycin (KAN), streptomycin (STR), amikacin (AMK), trimethoprim (TMP), chloramphenicol (CHL), tetracycline (TET), and sulfonamides (SUL). Plates were incubated at 37°C for 24 hours, and susceptibility was interpreted according to EUCAST clinical breakpoints (version 16.0).

### Genome analysis

Eighty-eight Hg^R^ isolates were sequenced using Illumina short-read technology, and 20 additional isolates were sequenced with Oxford Nanopore long-read technology to resolve chromosomal and plasmid synteny. Genomic DNA was extracted, quality-checked, and prepared for sequencing using standard Illumina or ONT library preparation workflows, and sequencing was performed on a HiSeq4000 (Illumina) or R10.4.1 flow cells (ONT) as recommended by manufacturers.

Quality control and read filtering were performed using fastp v0.24.0 ^67^. Paired-end reads were de novo assembled with SPAdes v4.0.0 ^68^, using isolate mode (isolate --) and a coverage cut-off of 100 (--cov-cutoff 100), excluding contigs with an average read depth below 100x. Long-read sequences were assembled with Unicycler v0.5.0. Assembly quality was assessed with QUAST v5.3.0, and long assemblies were polished with short reads using Pilon v1.24. Genome annotation was carried out using Bakta v1.10.3 ^69^ with a minimum contig length of 200 bp (--min-contig-length 200), excluding shorter contigs from annotation. Sequence types (STs) were assigned using the in silico MLST v2.23.0 tool (https://github.com/tseemann/mlst), and *E. coli* phylogroups were determined using EzClermont v0.7.0 (https://github.com/nickp60/EzClermont).

#### Resistome analysis

ARGs were identified with AMRFinderPlus v4.0.3 ^70^ using nucleotide sequence, while MRGs and biocide resistance genes (BRGs) were detected by querying the predicted protein sequences from each isolate against a custom database using BLASTp. This custom database was based on MRG included in Bacmet v2.0 ^71^ and supplemented with curated UniProt/Swiss-Prot sequences to improve the representation of allelic variants of MRGs already included in BacMet but not captured by the corresponding reference sequences. No additional MRG families or novel resistance genes were introduced. Detection thresholds were set at ≥80% sequence coverage and ≥90% identity for ARGs, and ≥80% sequence coverage and identity for MRGs and BRGs, as previously reported. Resistance profiles were summarised as presence–absence matrices and visualised as heatmaps using the R package *pheatmap*.

#### Operon and transposon analysis

The diversity of *mer*, *pco,* and *sil* operons was assessed by clustering sequences at an 85% amino-acid identity threshold using BLOSUM62-based pairwise alignments. MRGs were initially identified using thresholds of ≥80% coverage and ≥80% amino-acid identity against the metal-resistance database. A more stringent threshold of ≥85% amino-acid identity was then applied to identify closely related sequence variants within each MRG. This threshold was established based on sequence comparisons of individual Mer proteins, which showed high sequence conservation among closely related variants, whereas more divergent variants formed distinct groups. The same criterion was applied to *pco* and *sil* to ensure consistent classification across the three metal-resistance systems.

Each cluster was assigned a unique identifier within each gene. Operon synteny was visualised using the R package “gggenes”. For phylogenetic analysis, amino acid sequences were aligned gene by gene using Clustal Omega to ensure that only homologous sequences were compared, and the resulting alignments were concatenated according to the conserved gene order within each operon. When genes were absent, gaps corresponding to the length of the missing gene were introduced to preserve the operon structure. Phylogenetic trees were constructed for each operon using IQ-TREE, and trees were visualised and annotated with metadata using iTOL ^72^. *Mer* operons and associated transposons were classified using the TnCentral database ^73^. Putative transposons were identified by screening for key marker genes, including *tnpA* and *tnpR* (indicative of Tn*1696*-like elements) and *tnpA, tnpR,* and *tnpM* (indicative of Tn*21*-like elements). Transposase annotations were subsequently confirmed using TnCentral.

#### Plasmid analysis

Plasmids were characterised using three complementary and widely used approaches, COPLA^35^, replicon-typing ^36^, and network-based plasmid clustering, to capture different aspects of plasmid relatedness in the absence of a universal plasmid classification system.

Plasmids were reconstructed from FASTA assemblies, typed using the MOB-recon module of MOB-suite v3.1.9 ^36^, and annotated with Bakta^69^. A gene-by-gene presence-absence matrix was generated using the “accnet” function from the PATO R package, with a default similarity parameter of 70%. Pairwise Jaccard similarity was then calculated from this matrix. A k-nearest neighbor network (K-NNN) was then constructed with k = 10 and a Jaccard similarity threshold of 0.5, connecting each plasmid to its ten most similar counterparts ^74^. The value of *k* = 10 was selected following the approach implemented in PATO ^74^ and defines the maximum number of nearest neighbors considered for each plasmid, rather than the maximum number of nodes within a cluster. Plasmid clustering was initially conducted using mclust ^75^ and further refined using the Louvain community-detection algorithm implemented in *igraph*. The resulting network was visualised in Cytoscape and imported into R using the package *tidygraph*. This graph-based approach was used to define plasmid groups (PGs) based on sequence relatedness, providing a reference-independent classification of the plasmid sequences.

Plasmid features, including predicted mobility, replicase types, and relaxase IDs, were integrated into the network. Replicase types, MRGs, and ARGs were identified by submitting plasmid assemblies to MOB-suite, our custom MRG database, and the AMRFinder database, respectively ^70,71^. Plasmid sequence types were assigned using pMLST ^76^ while plasmid taxonomic units (PTUs) were determined with COPLA2 ^35^ (https://castillo.dicom.unican.es/copla2/), which assigns plasmids to taxonomic units based on genomic relatedness. Together with the PGs defined above, these approaches provided complementary views of plasmid relatedness and diversity.

All data manipulation and visualisation were performed using the *tidyverse* package. Within-group plasmid diversity was assessed using Clinker, using a minimum sequence identity threshold of 80%, and visualised as heatmaps, with closed plasmids serving as the reference for most PGs ^77^. Clinker analyses were performed using the plasmid sequences together with their Bakta gene annotations. Pairwise average nucleotide identity (ANI) between plasmids from the same PG and related plasmids from public databases was calculated using fastANI v.1.34 (https://github.com/ParBLiSS/FastANI). To identify similar previously characterised plasmids, sequences were queried against the Nucleotide BLAST database of the National Center for Biotechnology Information (NCBI).

### *Mer* operon occurrence in public databases

To assess the representation of MRGs across genomic datasets, isolates were compared with genomes from the AllTheBacteria database ^44^ through the BacTaxID web platform to examine the distribution of MRG across bacterial genera. Isolates were subsequently assigned to hierarchical bacterial typing groups using BacTaxID ^43^, and groups containing fewer than four isolates were excluded from comparative analysis. To ensure consistent annotation criteria across datasets, MRGs in our isolates were reannotated using AMRFinderPlus, with thresholds of ≥80% sequence coverage and ≥80% nucleotide identity to both our isolates and the AllTheBacteria genomes. Comparisons were based on representative marker genes for the corresponding resistance systems, including *mer*A, *pco*A, *sil*A, *ars*A, *iuc*A, *iro*B, and *ybt*P. Differences in the detection of these markers between datasets were assessed using Fisher’s exact test on 2 x 2 presence–absence contingency tables, with P values adjusted for multiple comparisons using the Benjamini-Hochberg procedure.

## Ethics approval and consent to participate

This study was conducted in accordance with the principles of the Declaration of Helsinki. Ethical approval was obtained from the Ethic Committees of the Hospital Universitari i Politècnic La Fe, Valencia (approval number 2012/0018), Parc de Salut MAR, Sabadell (approval number 2005/2016/I), and the Donostia Hospital, Guipuzkoa (approval number PI2014150). Written informed consent was obtained from the parents or legal guardians of all participants.

## DECLARATIONS

## Availability of data and material

All sequence data have been deposited in the European Nucleotide Archive (ENA) under project accession PRJEB115272. The datasets related to epidemiological data of individual children used to generate Figure 1 and Supplementary Figures 1 and 2 are not publicly available due to legal and ethical regulations. Nonetheless, they may be available upon request to the director of the INMA Project, Mónica Guxens in accordance with the local, national, and European Union regulations.

## Author’s contribution

C.L-L.: all bioinformatic and statistical analysis, including figures and tables; manuscript writing; A.R.: faecal sampling, bacterial processing, culturomics, manuscript revision; A.S.V.: faecal sampling, bacterial processing, manuscript revision; M-J.L-E., and S.L.P.: children recruitment (Valencia region), faecal collection, metadata collection, and manuscript revision; M.G. and M.B.: children recruitment (Sabadell region), faecal collection, metadata collection, and manuscript revision; A.I., and Z.B.: children recruitment (Gipuzkoa region), faecal collection, metadata collection, and manuscript revision; L.C.: children datasets for building Figure 1, and Supplementary Figures 1 and 2; F.B.: former collaborative study design, manuscript revision; V.F.L.: C.L-L’s bioinformatics training and supervision, bioinformatic and statistical design, data analysis, manuscript revision; T.M.C.: study design, data analysis, and manuscript writing. All the authors have read and approved the final version of this document.

## Acknowledgements

We thank Mercedes Rodríguez for technical support with faecal sample processing; Miguel Díaz Fernández de Bobadilla for supporting C.L-L. with initial bioinformatic analysis, Leticia Olavarrieta (UCA-IRYCIS) for sequencing support; and Susana Gros (IS Global) for coordinating the institutional collaboration.

Declaration of the Use of Generative AI and AI-Assisted Technologies in the Writing Process: During the preparation of this work, the authors used ChatGPT (OpenAI) and Grammarly to improve language and readability. The authors reviewed and edited the content as needed and took full responsibility for the original content of the publication.

## References

1. United Nations Environment Programme. Bracing for Superbugs: Strengthening environmental action in the One Health response to antimicrobial resistance (2023). http://www.unep.org/resources/superbugs/environmental-action

2. Coque, T. M., Cantón, R., Pérez-Cobas, A. E., Fernández-de-Bobadilla, M. D. & Baquero, F. Antimicrobial Resistance in the Global Health Network: Known Unknowns and Challenges for Efficient Responses in the 21st Century. Microorganisms 11, 1050 (2023).

3. Baker-Austin, C., Wright, M. S., Stepanauskas, R. & McArthur, J. V. Co-selection of antibiotic and metal resistance. Trends Microbiol. 14, 176–182 (2006).

4. Gillieatt, B. F. & Coleman, N. V. Unravelling the mechanisms of antibiotic and heavy metal resistance co-selection in environmental bacteria. FEMS Microbiol. Rev. 48, fuae017 (2024).

5. Pal, C. et al. Metal Resistance and Its Association With Antibiotic Resistance. Adv. Microb. Physiol. 70, 261–313 (2017).

6. Musah, B. I., Yang, J. & Xu, G. Drivers of toxic element accumulation in terrestrial ecosystems across elevational gradients. Ecol. Indic. 174, 113446 (2025).

7. Turner, R. J. The good, the bad, and the ugly of metals as antimicrobials. BioMetals 37, 545–559 (2024).

8. Grand View Research. Biocides Market Size, Share, Growth & Trends Report, 2030. Grand view research (2024). https://www.grandviewresearch.com/industry-analysis/biocides-industry

9. United Nations Environment Programme. Global Mercury Assessment 2018 (2019) https://www.unep.org/resources/publication/global-mercury-assessment-2018

10. Tchounwou, P. B., Yedjou, C. G., Patlolla, A. K. & Sutton, D. J. Heavy metal toxicity and the environment. Exp. Suppl. 2012 101, 133–164 (2012).

11. World Health Organization. Preventing disease through healthy environments: exposure to mercury: a major public health concern (2nd ed., 2021) https://www.who.int/publications/i/item/9789240023567

12. European Environment Agency. Mercury pollution remains a problem in Europe and globally (2018). https://www.eea.europa.eu/highlights/mercury-pollution-remains-a-problem

13. Zhao, M., Li, Y. & Wang, Z. Mercury and Mercury-Containing Preparations: History of Use, Clinical Applications, Pharmacology, Toxicology, and Pharmacokinetics in Traditional Chinese Medicine. Front. Pharmacol. 13, 807807 (2022).

14. Varkey, MI., Shetty, R. & Hegde, A. Mercury Exposure Levels in Children with Dental Amalgam Fillings. Int. J. Clin. Pediatr. Dent. 7, 180–185 (2014).

15. Porter, F. D., Silver, S., Ong, C. & Nakahara, H. Selection for mercurial resistance in hospital settings. Antimicrob. Agents Chemother. 22, 852–858 (1982).

16. Summers, A. O. et al. Mercury released from dental ‘silver’ fillings provokes an increase in mercury- and antibiotic-resistant bacteria in oral and intestinal floras of primates. Antimicrob. Agents Chemother. 37, 825–834 (1993).

17. Barkay, T., Miller, S. M. & Summers, A. O. Bacterial mercury resistance from atoms to ecosystems. FEMS Microbiol. Rev. 27, 355–384 (2003).

18. Christakis, C. A., Barkay, T. & Boyd, E. S. Expanded Diversity and Phylogeny of mer Genes Broadens Mercury Resistance Paradigms and Reveals an Origin for MerA Among Thermophilic Archaea. Front. Microbiol. 12, 682605 (2021).

19. Golysheva, A. A., Litvinenko, L. V. & Ivshina, I. B. Diversity of Mercury-Tolerant Microorganisms. Microorganisms 13, 1350 (2025).

20. World Health Organization. WHO bacterial priority pathogens list, 2024: Bacterial pathogens of public health importance to guide research, development and strategies to prevent and control antimicrobial resistance (2024). https://www.who.int/publications/i/item/9789240093461

21. Price, L. B., Hungate, B. A., Koch, B. J., Davis, G. S. & Liu, C. M. Colonizing opportunistic pathogens (COPs): The beasts in all of us. PLoS Pathog. 13, e1006369 (2017).

22. Castaño, A. et al. Mercury levels in blood, urine and hair in a nation-wide sample of Spanish adults. Sci. Total Environ. 670, 262–270 (2019).

23. Li, X. et al. Differential responses of the gut microbiome and resistome to antibiotic exposures in infants and adults. Nat. Commun. 14, 8526 (2023).

24. Llop, S. et al. Prenatal exposure to mercury and neuropsychological development in young children: the role of fish consumption. Int. J. Epidemiol. 46, 827–838 (2017).

25. López-González, U. et al. Exposure to mercury among Spanish adolescents: Eleven years of follow-up. Environ. Res. 231, 116204 (2023).

26. Ramon, R. et al. Prenatal mercury exposure in a multicenter cohort study in Spain. Environ. Int. 37, 597–604 (2011).

27. Soler-Blasco, R. et al. Exposure to mercury among 9-year-old Spanish children: Associated factors and trend throughout childhood. Environ. Int. 130, 104835 (2019).

28. Hobman, J. L., Essa, A. M. M. & Brown, N. L. Mercury resistance (mer) operons in enterobacteria. Biochem. Soc. Trans. 30, 719–722 (2002).

29. Essa, A. M. M., Julian, D. J., Kidd, S. P., Brown, N. L. & Hobman, J. L. Mercury Resistance Determinants Related to Tn21, Tn1696, and Tn5053 in Enterobacteria from the Preantibiotic Era. Antimicrob. Agents Chemother. 47, 1115–1119 (2003).

30. Martini, K. M., Boddu, S. S., Nemenman, I. & Vega, N. M. Maximum likelihood estimators for colony-forming units. Microbiol. Spectr. 12, e03946–23 (2024).

31. Herigstad, B., Hamilton, M. & Heersink, J. How to optimize the drop plate method for enumerating bacteria. J. Microbiol. Methods 44, 121–129 (2001).

32. Fernández-de-Bobadilla, M. D. et al. The antimicrobial gut resistome of the Wayampi reveals a shared background of antibiotic and metal resistance genes with industrialized populations, underscoring the “robust-yet-fragile” architecture of human gut microbiomes. Microbiome 14, 93 (2026).

33. Staehlin, B. M., Gibbons, J. G., Rokas, A., O’Halloran, T. V. & Slot, J. C. Evolution of a Heavy Metal Homeostasis/Resistance Island Reflects Increasing Copper Stress in Enterobacteria. Genome Biol. Evol. 8, 811–826 (2016).

34. Macesic, N. et al. Silent spread of mobile colistin resistance gene mcr-9.1 on IncHI2 ‘superplasmids’ in clinical carbapenem-resistant Enterobacterales. Clin. Microbiol. Infect. 27, 1856.e7–1856.e13 (2021).

35. Redondo-Salvo, S. et al. COPLA, a taxonomic classifier of plasmids. BMC Bioinformatics 22, 390 (2021).

36. Robertson, J. & Nash, J. H. E. MOB-suite: software tools for clustering, reconstruction and typing of plasmids from draft assemblies. Microb. Genomics 4, (2018).

37. Lipworth, S. et al. The plasmidome associated with Gram-negative bloodstream infections: A large-scale observational study using complete plasmid assemblies. Nat. Commun. 15, 1612 (2024).

38. Cazares, A. et al. Pre- and postantibiotic epoch: The historical spread of antimicrobial resistance. Science 390, eadr1522 (2025).

39. Curiao, T. et al. Association of Composite IS26-sul3 Elements with Highly Transmissible IncI1 Plasmids in Extended-Spectrum-β-Lactamase-Producing Escherichia coli Clones from Humans. Antimicrob. Agents Chemother. 55, 2451–2457 (2011).

40. Shaw, L. P. et al. The leading region of many plasmids is adapted for translational efficiency. Nucleic Acids Res. 54, gkag810 (2026).

41. Matlock, W. et al. Enterobacterales plasmid sharing amongst human bloodstream infections, livestock, wastewater, and waterway niches in Oxfordshire, UK. eLife 12, e85302 (2023).

42. Johnson, T. J. et al. Separate F-Type Plasmids Have Shaped the Evolution of the H30 Subclone of Escherichia coli Sequence Type 131. mSphere 1, e00121–16 (2016).

43. Fernández-de-Bobadilla, M. D. & Lanza, V. F. BacTaxID: A universal framework for standardized bacterial classification. bioRxiv (2025). 10.64898/2025.12.09.693184

44. Hunt, M., et al. AllTheBacteria: a community resource empowers biology and discovers novel peptide antibiotics. Preprint at 10.1101/2024.03.08.584059 (2026).

45. Skurnik, D. et al. Is exposure to mercury a driving force for the carriage of antibiotic resistance genes? J. Med. Microbiol. 59, 804–807 (2010).

46. Guo, G., Yumvihoze, E., Poulain, A. J. & Man Chan, H. Monomethylmercury degradation by the human gut microbiota is stimulated by protein amendments. J. Toxicol. Sci. 43, 717–725 (2018).

47. Zhu, Q. et al. Toxic and essential metals: metabolic interactions with the gut microbiota and health implications. Front. Nutr. 11, 1448388 (2024).

48. Becking, L. G. M. B. Geobiologie of inleiding tot de milieukunde. (W.P. Van Stockum & Zoon, 1934).

49. Wang, Y., Batra, A., Schulenburg, H. & Dagan, T. Gene sharing among plasmids and chromosomes reveals barriers for antibiotic resistance gene transfer. Philos. Trans. R. Soc. B Biol. Sci. 377, 20200467 (2021).

50. Letten, A. D., Hall, A. R. & Levine, J. M. Using ecological coexistence theory to understand antibiotic resistance and microbial competition. Nat. Ecol. Evol. 5, 431–441 (2021).

51. Zhu, S., Hong, J. & Wang, T. Horizontal gene transfer is predicted to overcome the diversity limit of competing microbial species. Nat. Commun. 15, 800 (2024).

52. Kottara, A., Carrilero, L., Harrison, E., Hall, J. P. J. & Brockhurst, M. A. The dilution effect limits plasmid horizontal transmission in multispecies bacterial communities. Microbiology 167, 001086 (2021).

53. Kerfoot, W. C., Harting, S. L., Rossmann, R. & Robbins, J. A. Elemental mercury in copper, silver and gold ores: an unexpected contribution to Lake Superior sediments with global implications. Geochem. Explor. Environ. Anal. 2, 185–202 (2002).

54. Mindlin, S. et al. Present-day mercury resistance transposons are common in bacteria preserved in permafrost grounds since the Upper Pleistocene. Res. Microbiol. 156, 994–1004 (2005).

55. Cain, A. K. & Hall, R. M. Evolution of IncHI2 plasmids via acquisition of transposons carrying antibiotic resistance determinants. J. Antimicrob. Chemother. 67, 1121–1127 (2012).

56. Hughes, V. M. & Datta, N. Conjugative plasmids in bacteria of the ‘pre-antibiotic’ era. Nature 302, 725–726 (1983).

57. Liebert, C. A., Hall, R. M. & Summers, A. O. Transposon Tn21, flagship of the floating genome. Microbiol. Mol. Biol. Rev. MMBR 63, 507–522 (1999).

58. Novais, A. et al. International spread and persistence of TEM-24 is caused by the confluence of highly penetrating enterobacteriaceae clones and an IncA/C2 plasmid containing Tn1696::Tn1 and IS5075-Tn21. Antimicrob. Agents Chemother. 54, 825–834 (2010).

59. Partridge, S. R. & Hall, R. M. The IS1111 family members IS4321 and IS5075 have subterminal inverted repeats and target the terminal inverted repeats of Tn21 family transposons. J. Bacteriol. 185, 6371–6384 (2003).

60. Nakahara, H., Ishikawa, T., Sarai, Y., Kondo, I. & Kozukue, H. Mercury resistance and R plasmids in Escherichia coli isolated from clinical lesions in Japan. Antimicrob. Agents Chemother. 11, 999–1003 (1977).

61. Wang, Y. & Dagan, T. The evolution of antibiotic resistance islands occurs within the framework of plasmid lineages. Nat. Commun. 15, 4555 (2024).

62. Fuhrman, J. A. Microbial community structure and its functional implications. Nature 459, 193–199 (2009).

63. Baquero, F. et al. Phenotypic copper resistance in aerobic intestinal bacteria from children with different levels of copper-exposure. J. Environ. Health Sci. 3, 1–13 (2017).

64. Clermont, O., Christenson, J. K., Daubié, A.-S., Gordon, D. M. & Denamur, E. Development of an allele-specific PCR for Escherichia coli B2 sub-typing, a rapid and easy to perform substitute of multilocus sequence typing. J. Microbiol. Methods 101, 24–27 (2014).

65. Johnson, J. R. et al. Rapid and specific detection, molecular epidemiology, and experimental virulence of the O16 subgroup within Escherichia coli sequence type 131. J. Clin. Microbiol. 52, 1358–1365 (2014).

66. European Committee on Antimicrobial Susceptibility Testing (EUCAST). Guidance Documents. https://www.eucast.org/bacteria/guidance-documents/

67. Chen, S., Zhou, Y., Chen, Y. & Gu, J. fastp: an ultra-fast all-in-one FASTQ preprocessor. Bioinformatics 34, i884–i890 (2018).

68. Bankevich, A. et al. SPAdes: A New Genome Assembly Algorithm and Its Applications to Single-Cell Sequencing. J. Comput. Biol. 19, 455–477 (2012).

69. Schwengers, O. et al. Bakta: rapid and standardized annotation of bacterial genomes via alignment-free sequence identification. Microb. Genomics 7, 000685 (2021).

70. Feldgarden, M. et al. AMRFinderPlus and the Reference Gene Catalog facilitate examination of the genomic links among antimicrobial resistance, stress response, and virulence. Sci. Rep. 11, 12728 (2021).

71. Pal, C., Bengtsson-Palme, J., Rensing, C., Kristiansson, E. & Larsson, D. G. J. BacMet: antibacterial biocide and metal resistance genes database. Nucleic Acids Res. 42, D737–743 (2014).

72. Letunic, I. & Bork, P. Interactive Tree Of Life (iTOL) v5: an online tool for phylogenetic tree display and annotation. Nucleic Acids Res. 49, W293–W296 (2021).

73. Ross, K. et al. TnCentral: a Prokaryotic Transposable Element Database and Web Portal for Transposon Analysis. mBio 12, e0206021 (2021).

74. Fernández-de-Bobadilla, M. D. et al. PATO: Pangenome Analysis Toolkit. Bioinforma. Oxf. Engl. 37, 4564–4566 (2021).

75. Fraley, C. & Raftery, A. Model-based methods of classification: using the mclust software in chemometrics. J. Stat. Softw. 18, (2007).

76. Jolley, K. A., Bray, J. E. & Maiden, M. C. J. Open-access bacterial population genomics: BIGSdb software, the PubMLST.org website and their applications. Wellcome Open Res. 3, 124 (2018).

77. Gilchrist, C. L. M. & Chooi, Y.-H. clinker & clustermap.js: automatic generation of gene cluster comparison figures. Bioinformatics 37, 2473–2475 (2021).

