## Supplementary information for "Bacterial Mercury Resistance Reveals a Robust Species-Structured Human Antimicrobial Mobilome"

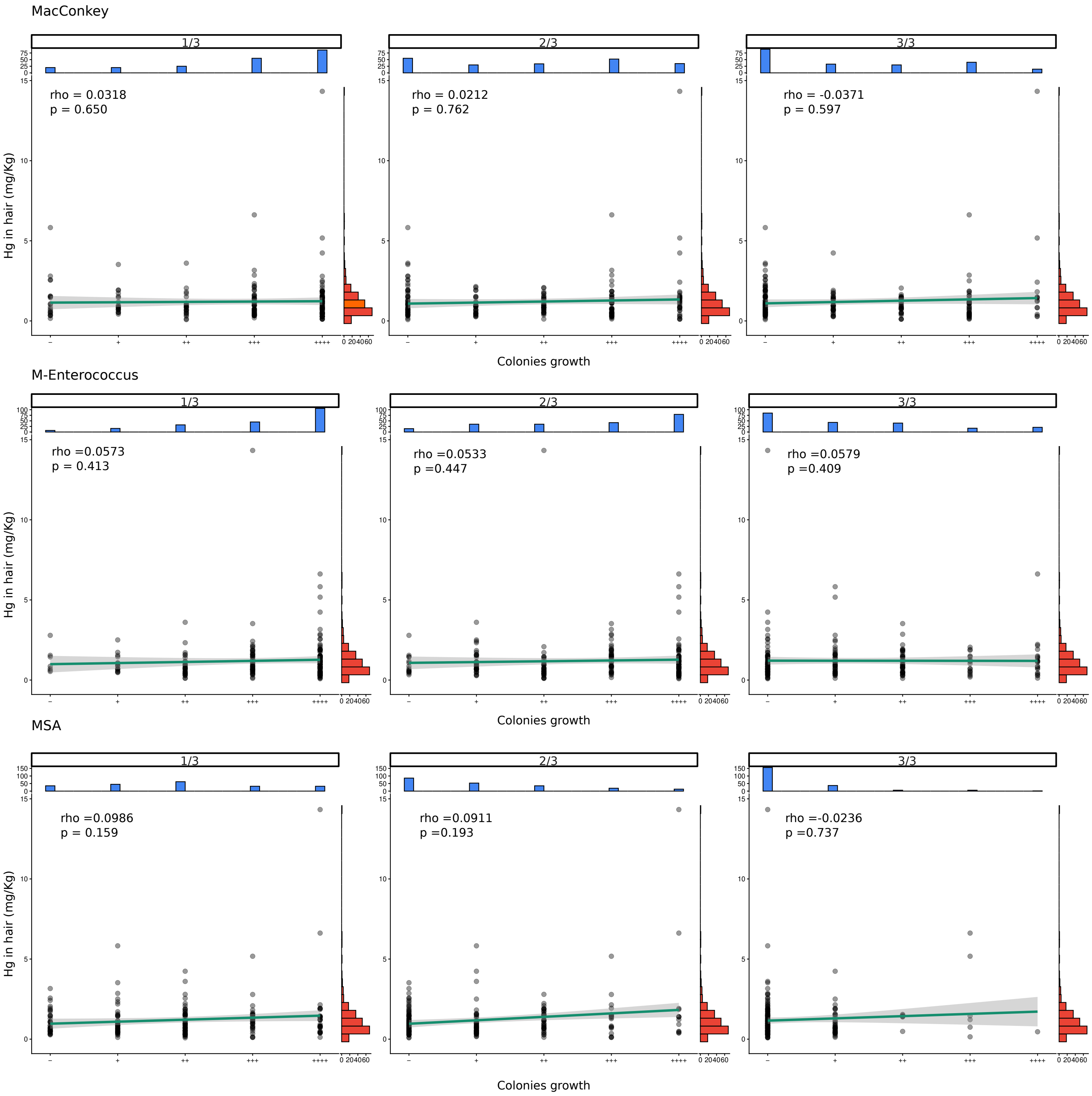


**Supplementary Figure 1.** Relationship between hair Hg concentrations and bacterial Hg tolerance. Hair Hg concentrations (mg·kg-1) measured at the 4-year visit were compared with Hg tolerance levels estimated from bacterial growth across Hg gradient agar plates: MacConkey, m-Enterococcus and mannitol salt agar (MSA). Susceptibility phenotypes were categorised by colony growth density as - (0 colonies), + (1-10), ++ (10-100), +++ (100-1000), and ++++ (>1000) across the 1/3, 2/3, and 3/3 gradient plate segments. Individual observations are shown together with fitted linear regression models and 95% confidence intervals. Marginal histograms (top) display the distribution of susceptibility categories, and side distributions summarise the distribution of hair Hg concentration. Spearman's correlation coefficients (rho) and corresponding P values are indicated in each panel. No significant association was observed between hair Hg concentrations and bacterial Hg tolerance.

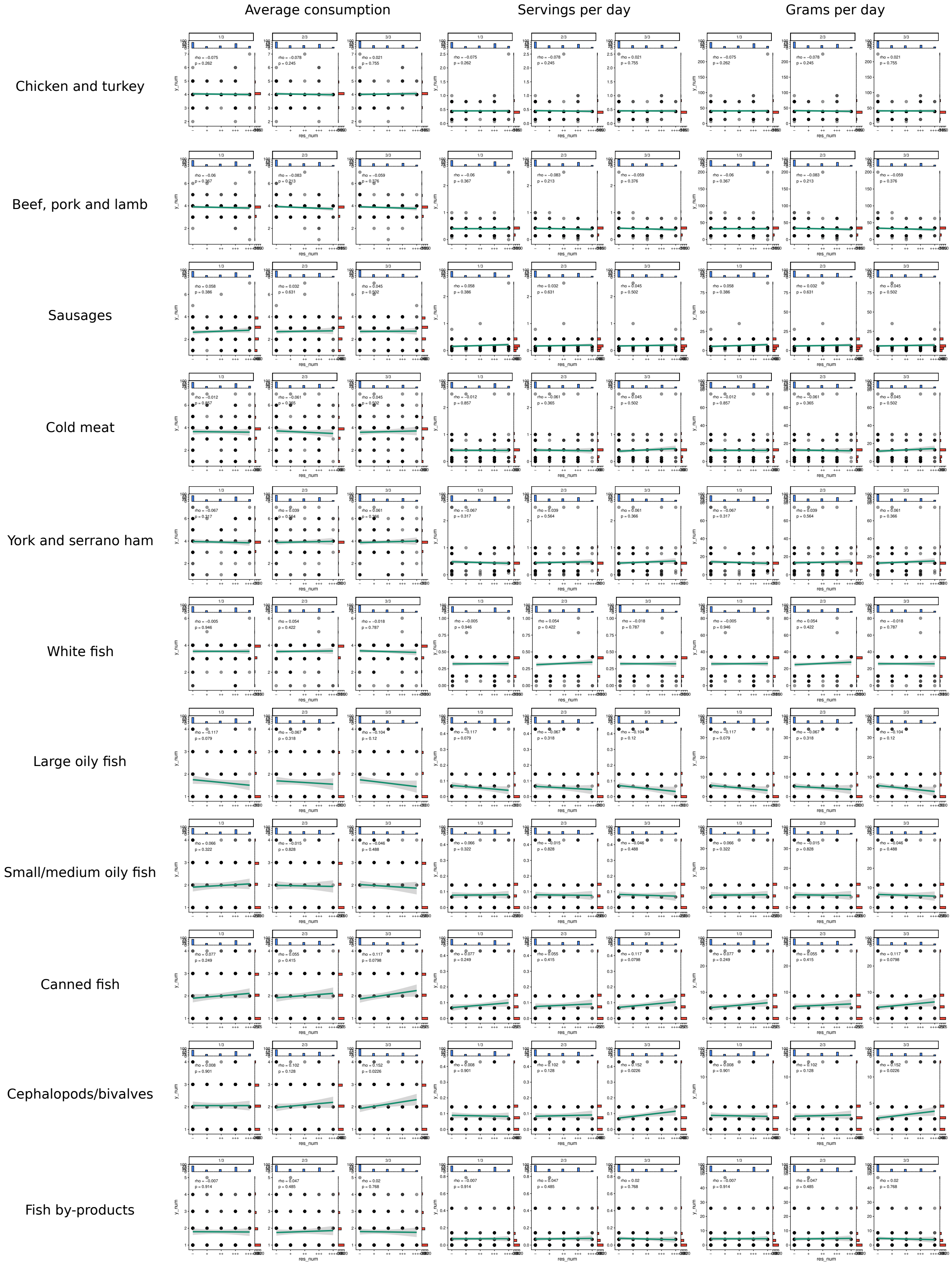


**Supplementary Figure 2.** Relationship between diet and bacterial Hg tolerance. Diet across 11 food groups, assessed using three dietary metrics: average consumption in the last year, servings per day, and grams per day, was compared with Hg susceptibility levels estimated from bacterial growth on MacConkey Hg gradient agar plates. Susceptibility phenotypes were categorised by colony growth density as - (0 colonies), + (1-10), ++ (10-100), +++ (100-1000), and ++++ (>1000) across the 1/3, 2/3, and 3/3 gradient plate segments. Individual observations are shown together with fitted linear regression models and 95% confidence intervals. Marginal histograms (top) display the distribution of susceptibility categories, and side distributions summarise the distribution of each dietary metric. Spearman's correlation coefficients (rho) and corresponding P values are indicated in each panel. No significant association was observed between the analysed dietary variables and bacterial Hg susceptibility across groups.

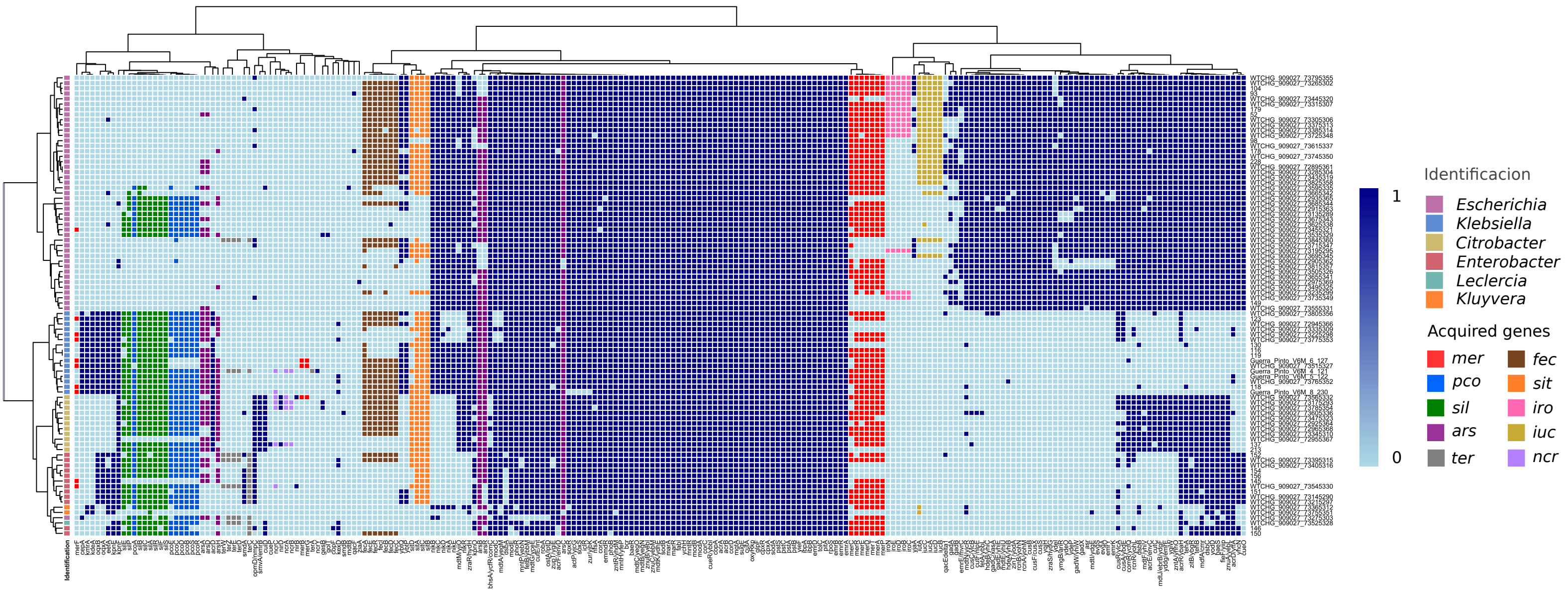


**Supplementary Figure 3.** The metalome of mercury-resistant Enterobacteriaceae. Heatmap showing the presence or absence of metal resistance genes (MRGs) and biocide resistance genes (BRGs) among Enterobacteriaceae in children, using 80% coverage and 80% identity thresholds for gene detection. Row annotation indicates the genus of each isolate. Different colours were used to indicate acquired MRGs (aMRGs), whereas intrinsic genes (iMRGs) are shown in blue. The aMRGs include *mer* (Hg), *pco* and *sil* (Cu), *ars* (As), *sit*, *iro*, *iuc* and *fec* (Fe), *ter* (Te) and *ncr* (Ni). All data are provided in Supplementary Table 2.

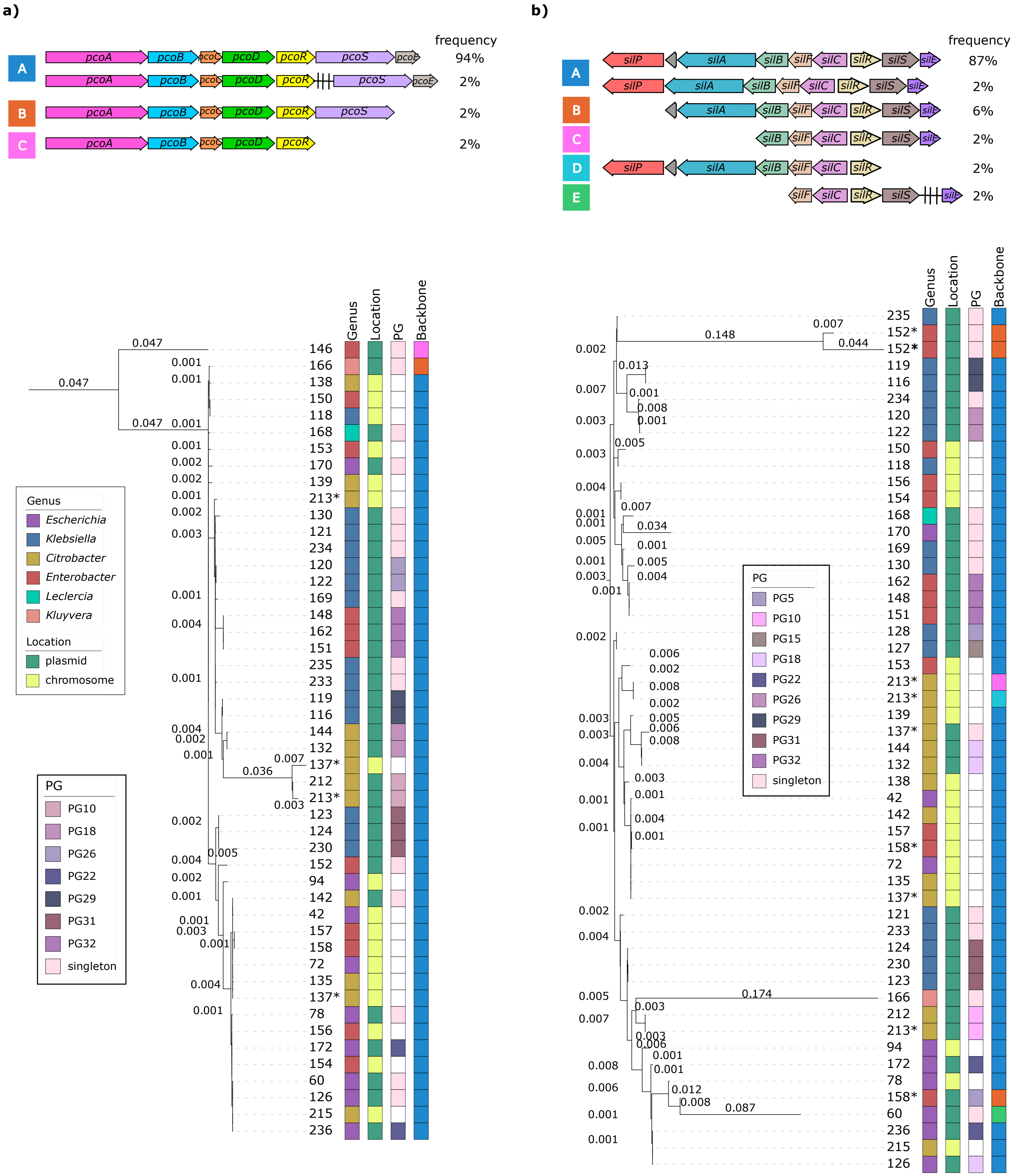


**Supplementary Figure 4.** Diversity of operons associated with acquired copper resistance. Schematic representations of the *pco* operon **(a)** and *sil* operon architectures **(b)** are shown. Arrows indicate gene content and orientation, and black bars represent intervening genes located between operon elements. Percentages denote the frequency of each configuration among *pco*- or *sil*-positive genomes. The accompanying phylogenetic tree highlights sequence diversity across isolates; asterisks indicate sequences derived from the same isolate. The phylogenetic tree was plotted and annotated using iTOL.

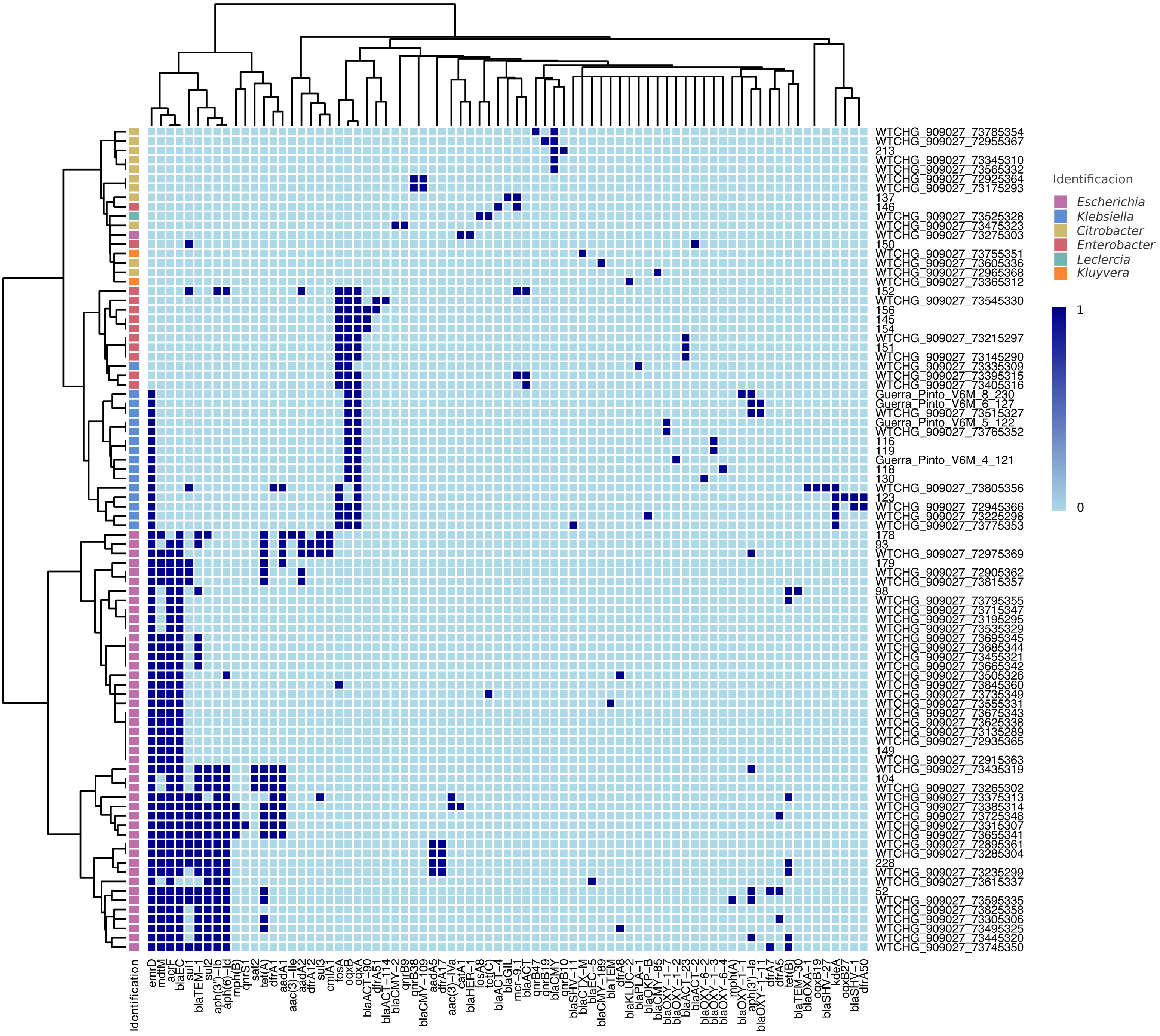


**Supplementary Figure 5.** The antibiotic resistome of mercury-resistant Enterobacteriaceae. Heatmap showing the presence or absence of antibiotic resistance genes (ARGs) among Enterobacteriaceae in children, using 80% coverage and 90% identity thresholds for gene detection. Row annotation indicates the genus of each isolate. All data are provided in Supplementary Table 3.

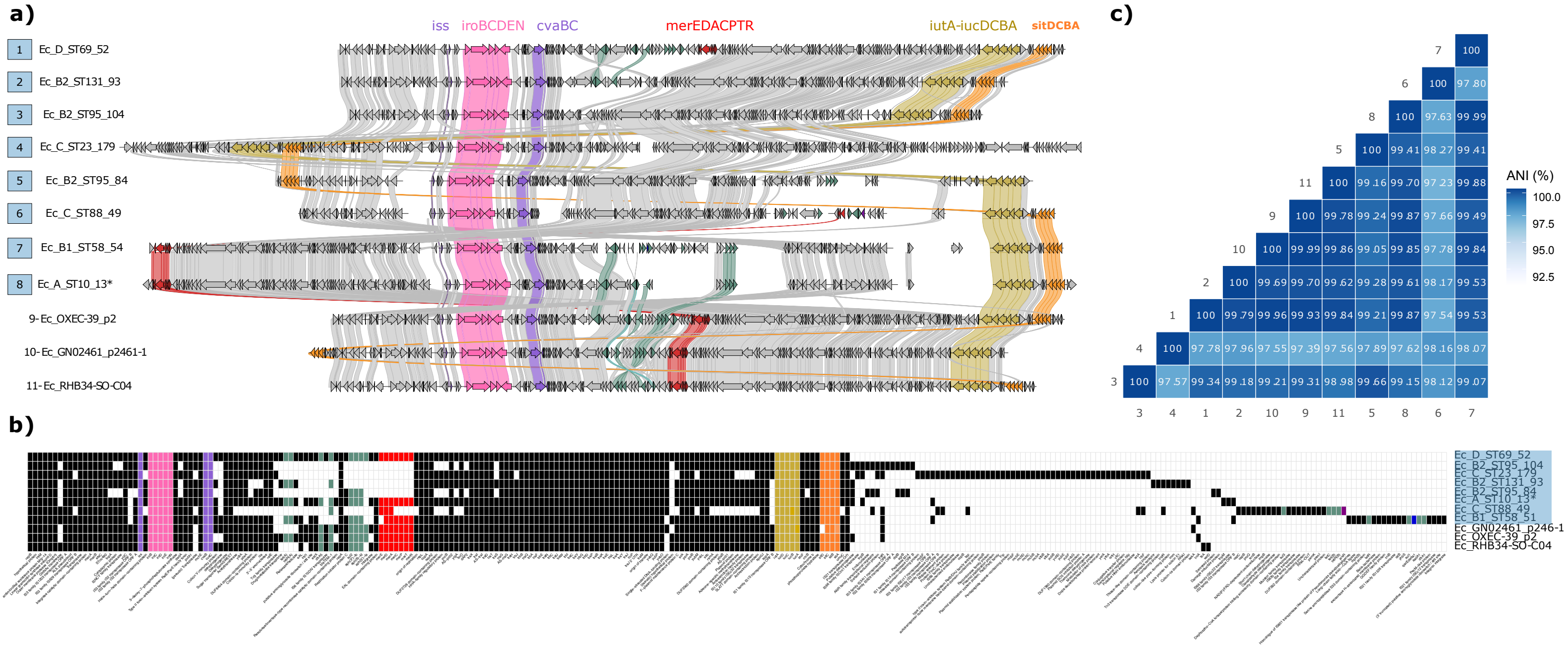


**Supplementary Figure 6**. Synteny and gene-content analysis of plasmids belonging to PG1. Plasmid sequences from this study and closely related plasmids retrieved from public databases were included in the analysis. (a) Structural comparison of representative plasmids using clinker. MRGs were annotated using Bakta. Isolates 1 and 2 correspond to closed long-read plasmids. (b) Gene presence–absence matrix, using the first isolate (designed as "isolate 1") as the reference. (a,b) Asterisks indicate transposons (Tn). Different colours denote acquired MRGs (as labelled in a), ARGs (green), and virulence factors (lilac). Sequences generated in this study are highlighted with blue boxes. (c) ANI matrix of the plasmids. Database plasmids showed 100% coverage and 99.99% identity (isolate 9), 98% coverage and 99.98% identity (isolate 10), and 97% coverage and 100% identity (isolate 11) relative to the reference plasmid.

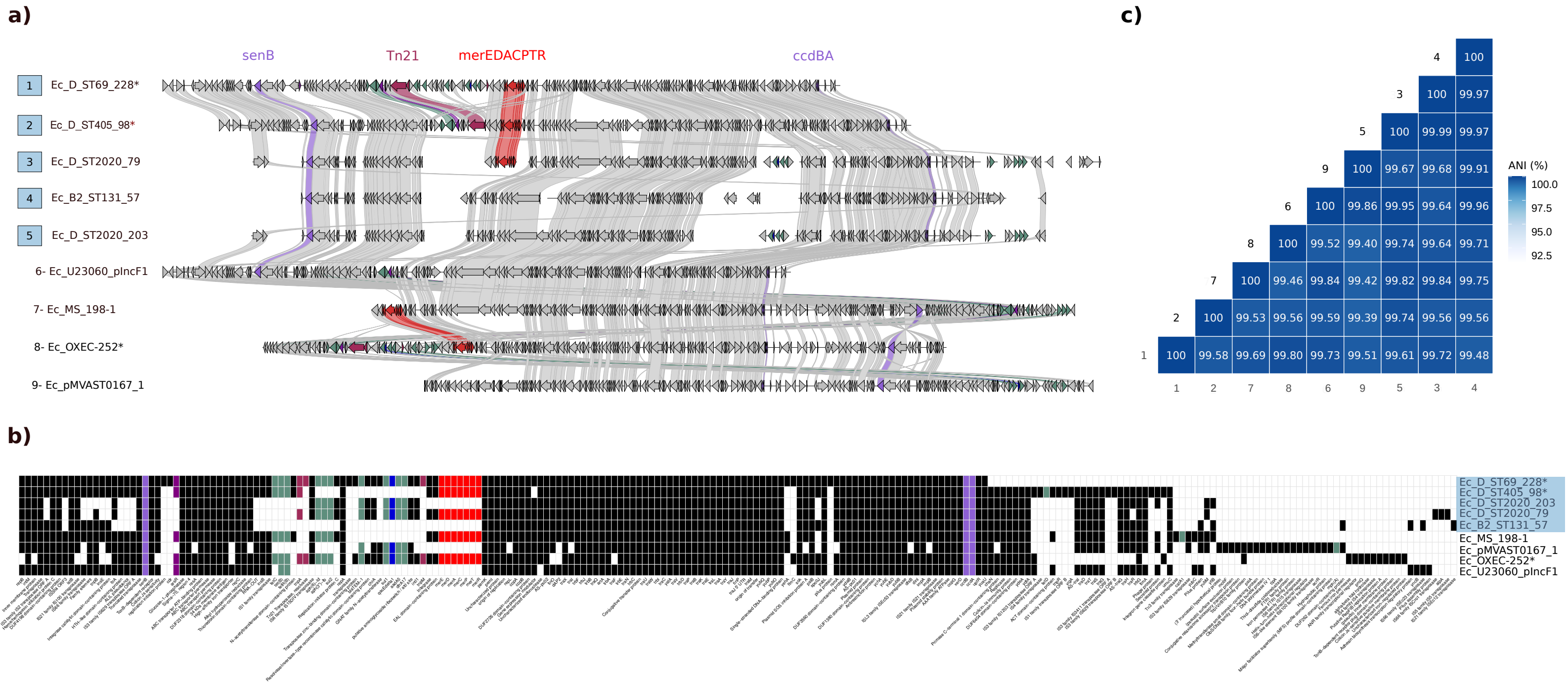


**Supplementary Figure 7.** Synteny and gene-content analysis of plasmids belonging to PG28. See legend Supplementary Figure 6. Isolates 1 and 2 correspond to closed long-read plasmids. Database plasmids showed 77% coverage and 99.95% identity (isolate 6), 89% coverage and 99.95% identity (isolate 7), 96% coverage and 99.96% identity (isolate 8), and 82% coverage and 99.91% identity (isolate 9) relative to the reference plasmid.

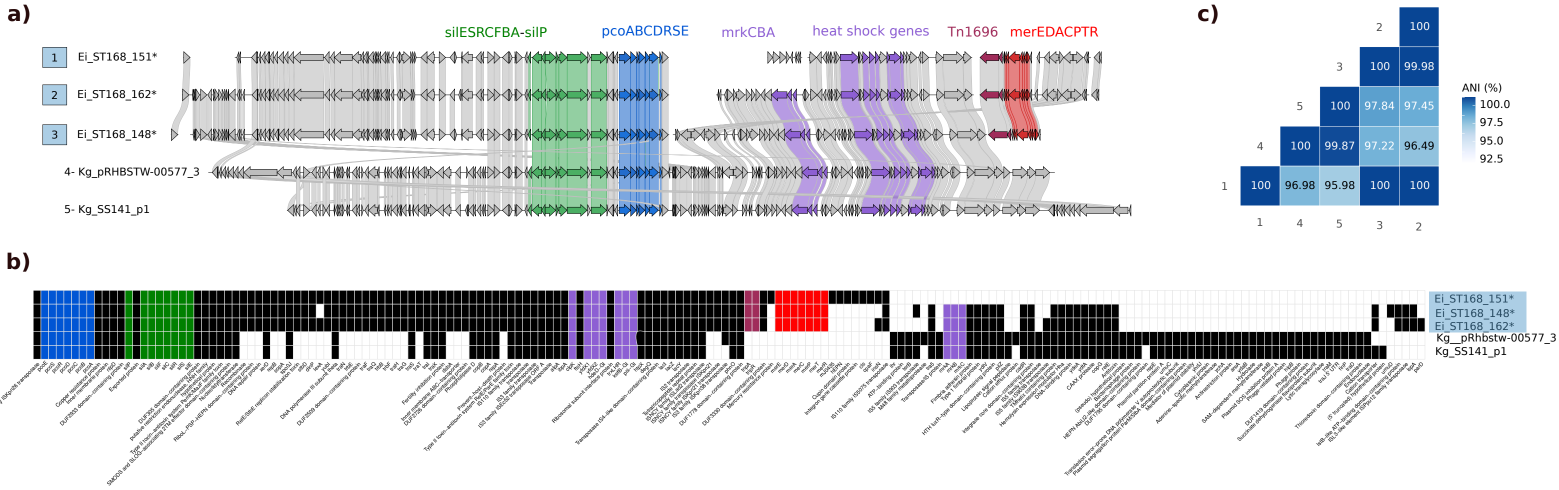


**Supplementary Figure 8.** Synteny and gene-content analysis of plasmids belonging to PG32. See legend Supplementary Figure 6. Database plasmids showed 59% coverage and 99.99% identity (isolate 4), and 58% coverage and 99.99% identity (isolate 5) relative to the reference plasmid. **(a)** Heat shock genes refer to *clpK*, *yfdX1*, *yfdX2*, *hde*D-GI, *trxLHR*, *kefB*-GI, *psi*-GI, all located within the heat resistance locus (LHR), a genomic island.

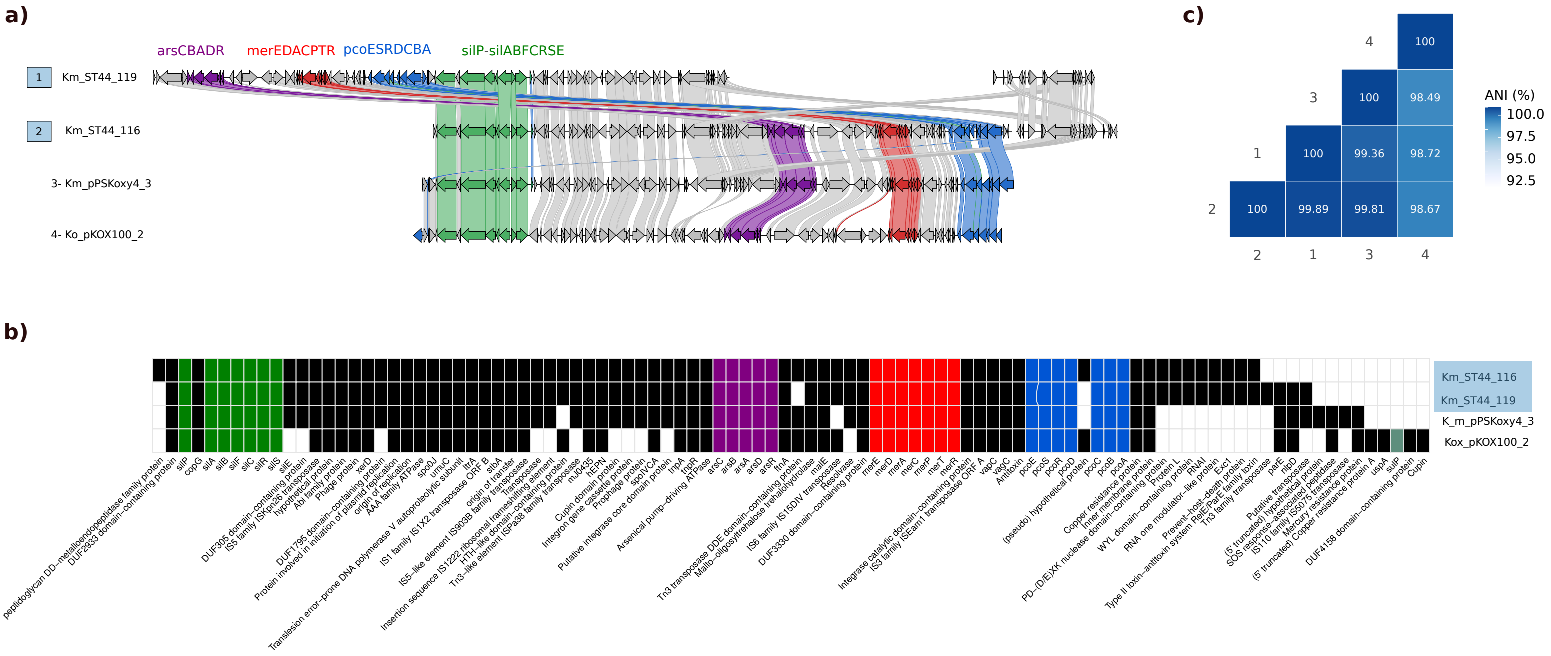


**Supplementary Figure 9.** Synteny and gene-content analysis of plasmids belonging to PG29. See legend Supplementary Figure 6. Isolates 1 and 2 correspond to closed long-read plasmids. Database plasmids showed 100% coverage and 100% identity (isolate 3), and 92% coverage and 100% identity (isolate 4) relative to the reference plasmid.

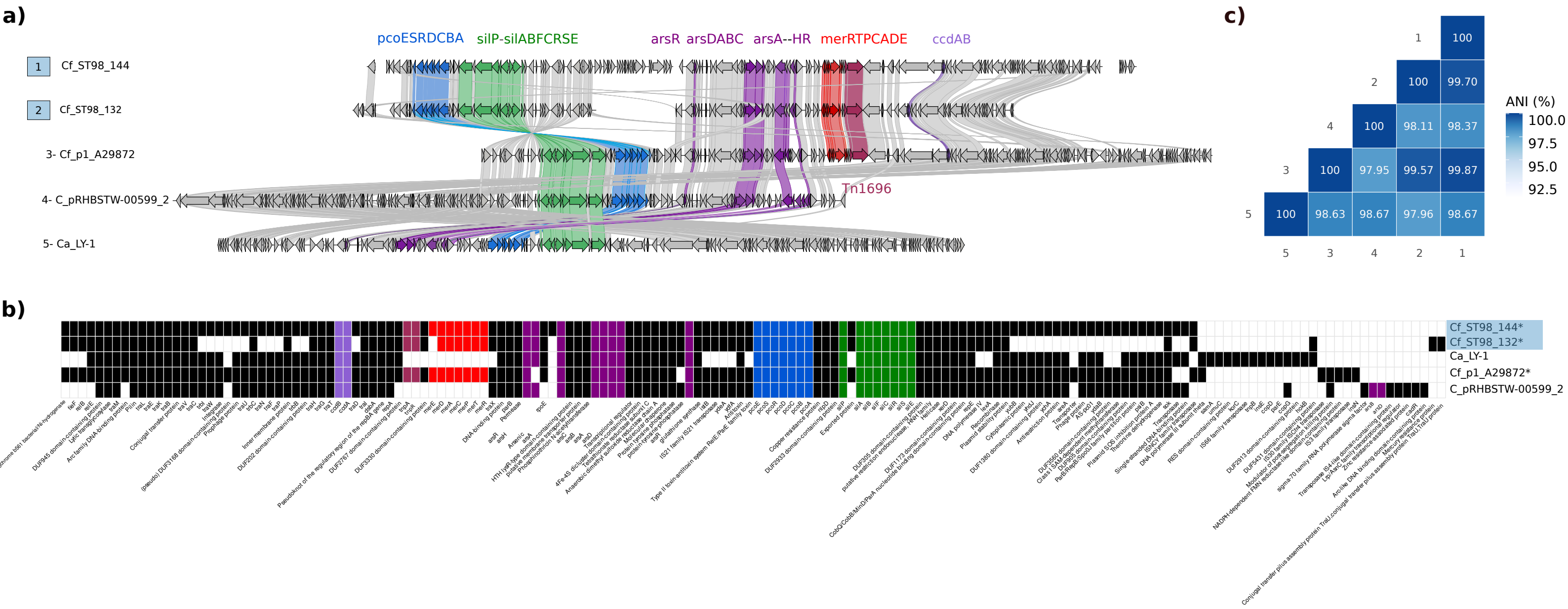


**Supplementary Figure 10.** Synteny and gene-content analysis of plasmids belonging to PG18. See legend Supplementary Figure 6. Database plasmids showed 98% coverage and 99.98% identity (isolate 3), 85% coverage and 97.85% identity (isolate 4), and 87% coverage and 99.85% identity (isolate 5) relative to the reference plasmid.

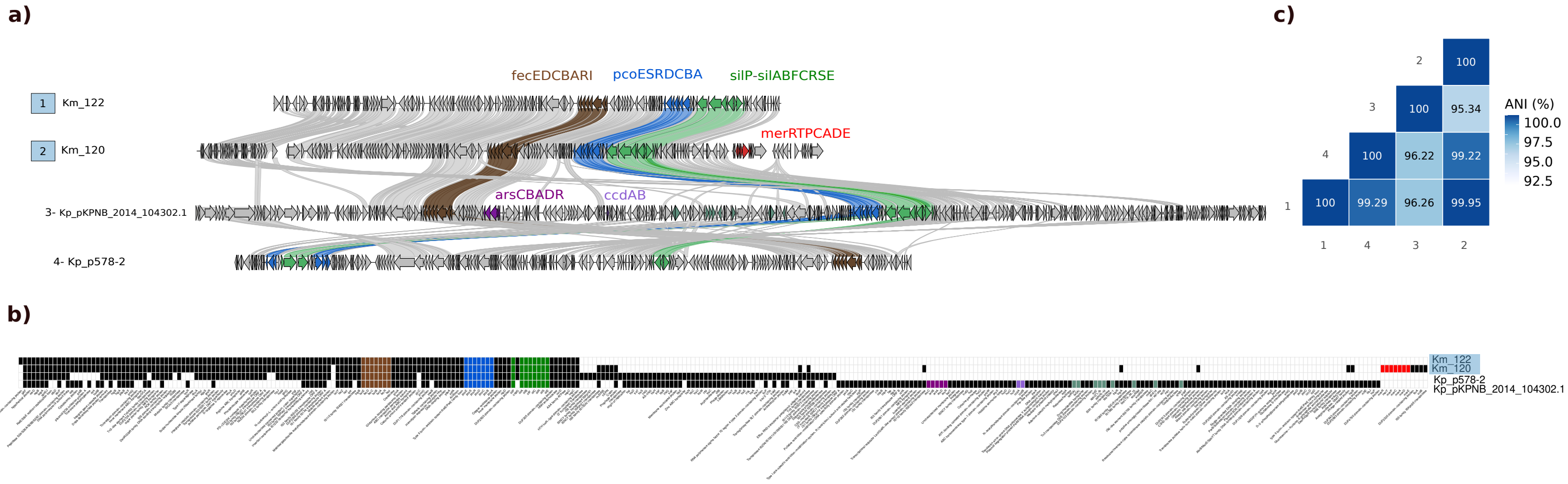


**Supplementary Figure 11.** Synteny and gene-content analysis of plasmids belonging to PG26. See legend Supplementary Figure 6. Isolate 1 corresponds to a closed long-read plasmids. Database plasmids showed 70% coverage and 97.76% identity (isolate 3), and 92% coverage and 99.81% identity (isolate 4) relative to the reference plasmid.

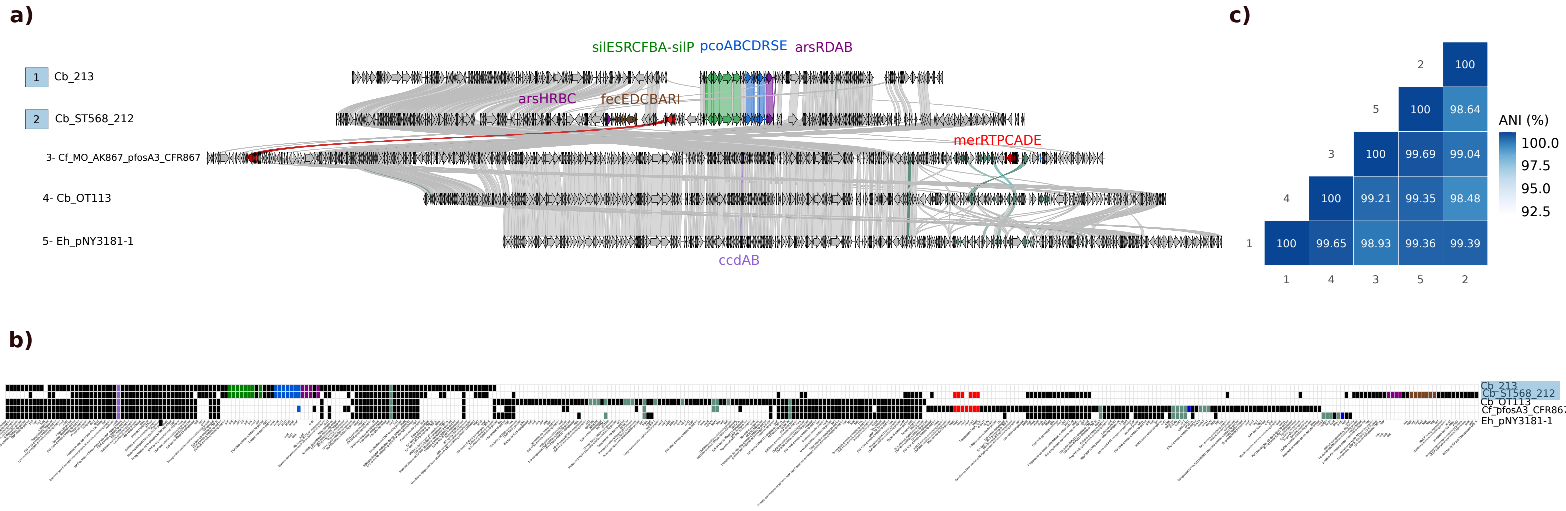


**Supplementary Figure 12.** Synteny and gene-content analysis of plasmids belonging to PG10. See legend Supplementary Figure 6. Isolate 1 corresponds to a closed long-read plasmids. Database plasmids showed 76% coverage and 99.98% identity (isolate 3), 73% coverage and 99.98% identity (isolate 4), and 73% coverage and 99.98% identity (isolate 5) relative to the reference plasmid.

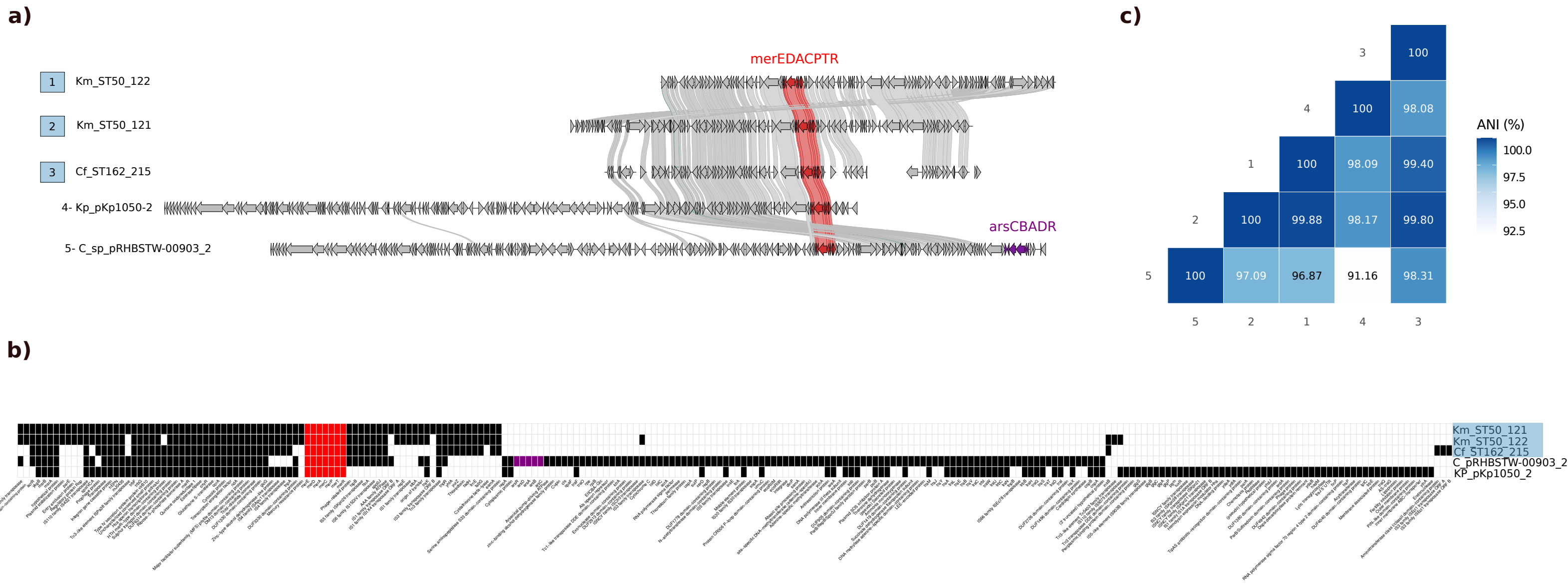


**Supplementary Figure 13.** Synteny and gene-content analysis of plasmids belonging to PG30. See legend Supplementary Figure 6. Isolates 1 and 2 correspond to closed long-read plasmids. Database plasmids showed 63% coverage and 99.94% identity (isolate 4), and 56% coverage and 99.89% identity (isolate 5) relative to the reference plasmid.

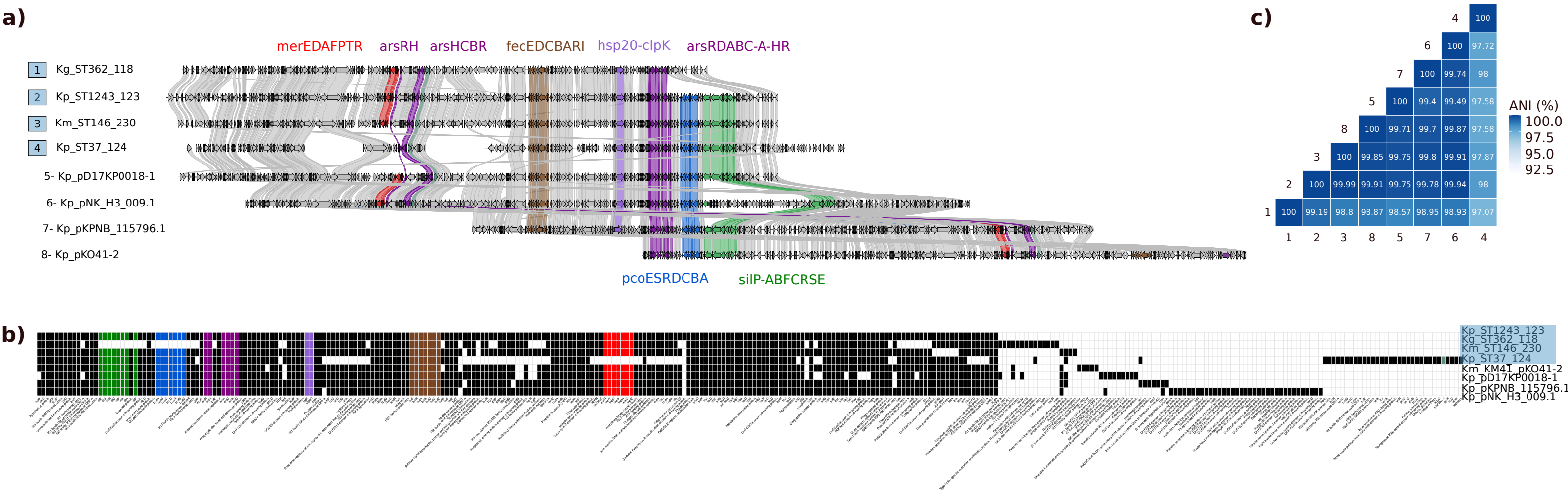


**Supplementary Figure 14.** Synteny and gene-content analysis of plasmids belonging to PG31. See legend Supplementary Figure 6. Isolates 1, 2 and 3 correspond to closed long-read plasmids. Database plasmids showed 95% coverage and 99.98% identity (isolate 5), 100% coverage and 99.99% identity (isolate 6), 100% coverage and 99.87% identity (isolate 7), and 97% coverage and 99.98% identity (isolate 8) relative to the reference plasmid.

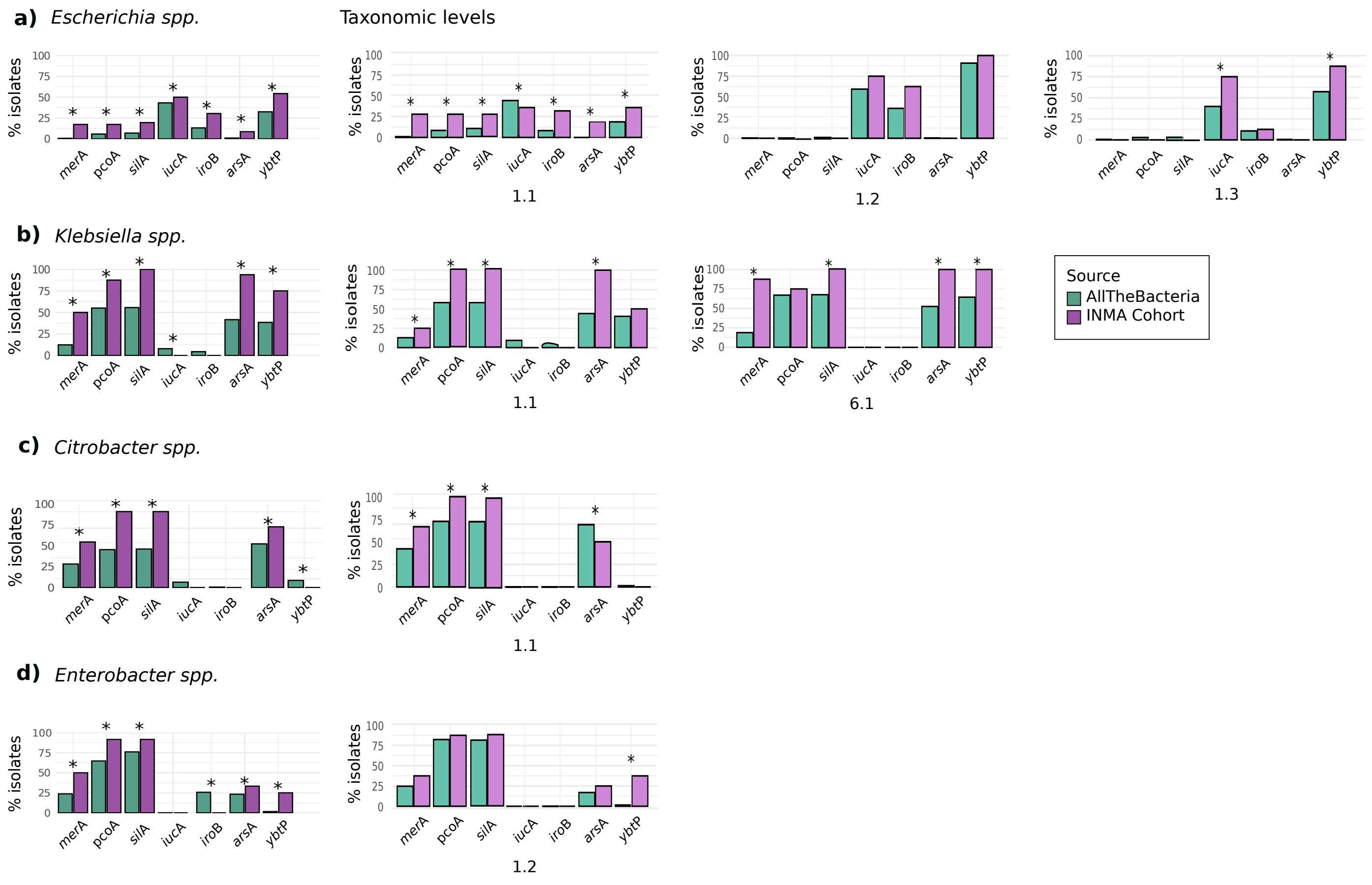


**Supplementary Figure 15.** Distribution of MRGs in databases. Comparison of the presence of MRGs in our isolates and in the AllTheBacteria database (https://allthebacteria.org/) across different genera: *Escherichia* **(a)**, *Klebsiella* **(b)**, *Citrobacter* **(c)** and *Enterobacter* **(d)**. Darker colours indicate genus-level comparison, whereas lighter colours indicate BactaxID-level representatives for each genus. Asterisks indicate significant differences (Supplementary Table 8).

**Supplementary Data 1. Epidemiological characteristics of mercury-resistant Enterobacterales isolates.** *mer*A indicates PCR-positive isolates, *merA*-/hybridisation+ indicates isolates negative by PCR but positive by hybridisation and *merA*-/hybridisation- indicates isolates negative by both methods. PhG denotes the phylogenetic groups for *E. coli* strains and sub-typing of B2 isolates, whereas NT indicates non-typable strains. Samples were collected from Sabadell (S), Gipuzkoa (G), and Valencia (V).

**Supplementary Data 2. Metal and biocide resistance genes identified using a curated database.** PhG denotes phylogenetic groups for *E. coli* strains; ST denotes sequence type by using MLST schemes (https://github.com/tseemann/mlst).

**Supplementary Data 3. Antimicrobial resistance genes identified using AMRfinderplus.** Abbreviations are provided in **Supplementary Table 2**.

**Supplementary Data 4. Plasmidome features of Hg-resistant isolates identified using MOB-suite.** PG denotes plasmid group; additional abbreviations are provided in **Supplementary Table 2**.

**Supplementary Data 5. Mercury-resistant plasmid groups.** PG denotes plasmid group; pMLST denotes the plasmid sequence type [76]; ARGs denotes antibiotic resistance genes; PhG denotes the phylogenetic group of *E. coli* strains; ST denotes sequence type using MLST schemes (https://github.com/tseemann/mlst); and PTU denotes plasmid taxonomic unit [35] (https://castillo.dicom.unican.es/copla2/).

**Supplementary Data 6. Mercury-resistant plasmid singletons.** Abbreviations are provided in **Supplementary Table 5**.

**Supplementary Data 7. Genomic location of MRGs in HgR Enterobacterales.** Green cells indicate elements carrying *mer* operons or MRGs co-localised within the same genetic element. Brown cells indicate siderophores and purple cells indicate other MRGs (*pco*, *sil*, *fec* and *ars*). Lighter colour denotes chromosomal (chr) location. Boxed cells indicate MRGs located on different elements, and bold text indicates MRGs present in more than one element. Abbreviations are provided in **Supplementary Table 5.**

**Supplementary Data 8. Statistical comparison of MRGs prevalence between isolates from the INMA cohort and AllTheBacteria databases.** Code denotes genus-level classification indicated as “General”, whereas numeric codes represent taxonomic levels according to BactaxID [43].
